# 53BP1/RIF1 and DNA-PKcs show distinct genetic interactions with diverse chromosomal break repair outcomes

**DOI:** 10.1101/2025.05.08.652920

**Authors:** Kaela Makins, Metztli Cisneros-Aguirre, Felicia Wednesday Lopezcolorado, Jeremy M. Stark

## Abstract

DNA double strand breaks (DSBs) are the effective lesion of cancer radiotherapy and induce gene editing. 53BP1 accumulates at DSBs and is implicated in end joining (EJ) repair, but its influence on DSB repair is distinct from canonical non-homologous end joining (C-NHEJ). We sought to define the genetic interplay of 53BP1 with C-NHEJ, focusing on the DNA-PKcs kinase. We examined Cas9 DSBs, which largely generates blunt DSBs, since blunt DSB EJ is dependent on C-NHEJ. Loss of 53BP1 does not affect blunt DSB EJ, but causes a reduction in such repair in DNA-PKcs deficient cells. In contrast, disrupting 53BP1 and DNA-PKcs, both alone and together, has similar effects on the type of deletion mutation (increase in microhomology deletions). We found similar effects on EJ with RIF1 loss, which is a downstream effector of 53BP1. Thus, 53BP1/RIF1 appear to play a backup role for DNA-PKcs during blunt DSB EJ, but function in the same pathway to suppress microhomology deletions. In contrast, 53BP1 and DNA-PKcs function independently to suppress homology-directed repair. Finally, DNA-PKcs kinase inhibition causes marked radiosensitivity, which is not additive with loss of 53BP1 and RIF1. Altogether, 53BP1/RIF1 and DNA-PKcs show distinct genetic interactions with diverse DSB repair outcomes.

## INTRODUCTION

Chromosomal DNA double strand breaks can be caused by physiological processes, as well as clastogenic cancer therapeutics (e.g. radiotherapy), or with targeted nucleases for gene editing. DSBs can be repaired by a number of pathways^1,2^. For one, homology-directed repair (HDR) uses invasion and replication of a template (e.g., the sister chromatid) to bridge the DSB. In contrast, end joining (EJ) repair involves ligation of the DSB ends without invasion of a homologous template. There are several sub-pathways of EJ repair that differ by the factors involved, the possible substrates for ligation, and genetic outcomes. Canonical non-homologous end joining (C-NHEJ) involves DNA-PKcs, Ku70/Ku80, XRCC4 and XLF, which synapse DSB ends, regulate DSB end processing, and position DNA Ligase 4 (LIG4) to catalyze the ligation^3–8^. C-NHEJ is critical for DSB repair events that are not stabilized by an annealing intermediate (e.g., blunt DSB ends), as well as EJ events involved in antibody maturation (i.e., V(D)J and Class Switch Recombination)^3–6,9,10^. In the absence of C-NHEJ factors, EJ remains relatively robust, but the outcomes show greater microhomology at the junctions, indicating that microhomology annealing is involved in bridging the DSBs^11–15^. Such annealing appears to be mediated by DNA polymerase theta (POLθ) although DNA polymerase lambda appears to also play a role^16,17^.

A key DNA damage response (DDR) factor involved in regulation of DSB repair is 53BP1, which was initially identified in a two-hybrid screen for interacting proteins with the p53 tumor suppressor^18–20^. Subsequent characterization found that 53BP1 can form robust foci at DSBs in cells, and can bind DNA and enhance DNA ligase 4 activity^18–20^. Additional evidence that 53BP1 is involved in DSB repair, and EJ in particular, is its key role in class switch recombination, and fusion of deprotected telomere ends^21,22^. However, 53BP1 has relatively modest effects on V(D)J recombination, which is dependent on core C-NHEJ factors (e.g., XRCC4)^23,24^. Another key distinction between 53BP1 and C-NHEJ is their influence on homology-directed repair (HDR). Namely, 53BP1 is critical to suppress HDR in BRCA1 deficient cells, i.e., loss of 53BP1 rescues HDR in BRCA1 deficient cells^25,26^. In contrast, loss of C-NHEJ factors (e.g., LIG4) do not appear to cause such rescue^25^. To suppress such HDR, 53BP1 appears to block DSB end resection in BRCA1 deficient cells^25,26^.

There are several other distinctions between 53BP1 and C-NHEJ. Whereas the C-NHEJ complex is centered around the KU heterodimer binding DNA ends^3–7^, 53BP1 appears to associate with DSBs by binding to specific chromatin marks adjacent to DSBs^27^. Such chromatin binding by 53BP1 promotes DSB recruitment of RIF1, a factor that binds phosphorylated 53BP1^27–38^. In vertebrates, RIF1 is critical for the timing regulation of DNA replication initiation and is a key mediator for 53BP1-dependent DSB repair^27–39^. Regarding the latter function, RIF1 bridges the REV7-Sheildin complex to 53BP1-bound chromatin at DSB sites, which is important to suppress DSB end resection^27–38^. In addition to suppressing end resection via Shieldin, 53BP1 has also been implicated in DSB end tethering and mobility at long distances^21–24^, as well as compaction of chromatin at DSBs^40^. These latter functions of 53BP1 may be linked to its capacity to oligomerize and form large condensates^41^, as well as form microdomains along with RIF1^40^. In summary, while 53BP1 and its effector RIF1 are distinct from C-NHEJ, and the interplay between these pathways is poorly understood. Thus, we sought to examine the genetic relationships between 53BP1/RIF1 and C-NHEJ on diverse DSB repair outcomes, focusing on the DNA-PKcs kinase subunit of C-NHEJ.

## RESULTS

### Combining loss of 53BP1 with an inhibitor of DNA-PKcs causes a decrease in No Indel EJ

We have sought to define the influence of 53BP1 on chromosomal break end joining, including whether this factor is partially redundant with C-NHEJ factors. To begin with, we examined the role of 53BP1 in blunt DSB EJ, which is dependent on several C-NHEJ factors^9^. Specifically, we used HEK293 cells with the chromosomal EJ7-GFP reporter, which measures EJ between two Cas9-induced blunt DSBs that are ligated without inserted or deleted nucleotides (nt) (No Indel EJ, Figure 1A)^9^. This No Indel EJ repair outcome yields ligation of the two blunt DSB ends, restoring GFP+ expression in cells, which can be measured via flow cytometry. This repair event is dependent the C-NHEJ factors XLF and KU, but only partially dependent on DNA-PKcs^42^.

**Figure 1.**
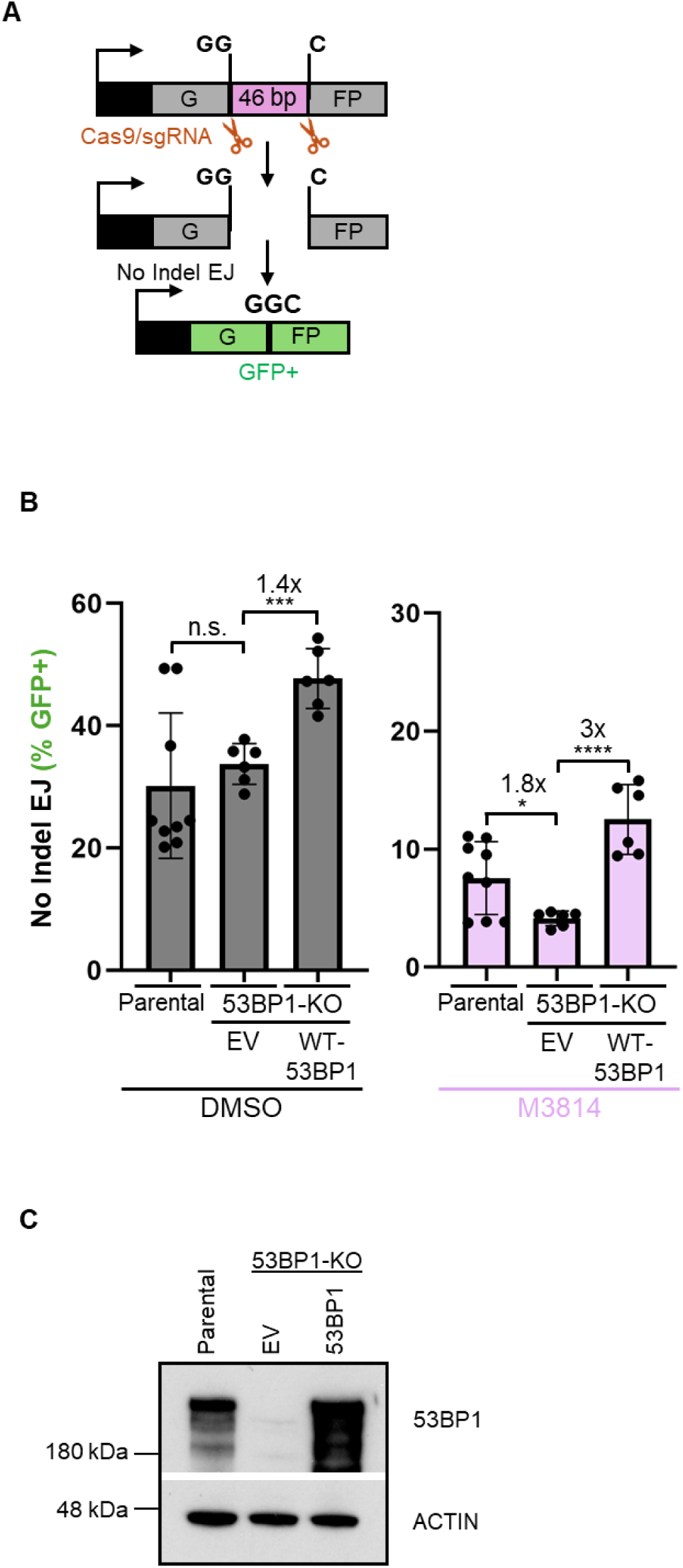
Combining loss of 53BP1 with an inhibitor of DNA-PKcs causes a decrease in No Indel EJ. **(A)** Schematic of the EJ7-GFP reporter assay, which is chromosomally integrated in HEK293 cells and measures No Indel EJ (i.e., repair of blunt DSB distal ends of two Cas9 DSBs). Cells are transfected with Cas9 and two sgRNA expression vectors, and GFP frequencies are normalized to transfection efficiency with parallel GFP transfections. **(B)** 53BP1 is not important for No Indel EJ, but becomes important with inhibition of DNA-PKcs. n=6, except Parental cells n=9. Statistics with unpaired t-test using Holm-Sidak correction. *P<0.5, ***P<0.001, ****P<0.0001, n.s.=not significant. **(C)** Immunoblot analysis of 53BP1 for the cell lines and conditions shown in (B).

We considered that 53BP1 may be partially redundant with DNA-PKcs for No Indel EJ. We based this hypothesis on evidence that both of these factors are implicated in DSB end synapsis, and regulation of DSB end processing^3–8,21–26,40^. Using the EJ7-GFP reporter integrated into HEK293 cells (Parental), we generated a 53BP1 knockout (53BP1-KO) cell line, and examined the frequency of this repair event, both with and without treatment with a DNA-PKcs kinase inhibitor (M3814, 500 nM)^43,44^. We also included transient expression of 53BP1 to test for complementation. We found no significant difference in No Indel EJ frequencies between Parental and 53BP1-KO cells, and expression of 53BP1 in the 53BP1-KO caused a modest increase (1.4-fold, Figure 1B). These findings are similar to prior studies with 53BP1-KO mouse embryonic stem cells (mESCs)^45^. However, M3814 treatment caused a reduction in No Indel EJ that was further reduced with 53BP1 loss (1.8-fold), which was complemented with expression of 53BP1 (3-fold). As a control, we found that 53BP1 loss does not affect M3814 inhibition of DNA-PKcs kinase activity (Supplementary Figure 1). These findings indicate that loss of 53BP1 alone has no obvious effect on No Indel EJ, but when combined with DNA-PKcs inhibition, loss of 53BP1 causes a decrease in this EJ event.

### Combining loss of 53BP1 with either a DNA-PKcs inhibitor or genetic loss causes a decrease in DSB blunt EJ

We then sought to use a distinct assay to examine the interplay between 53BP1 and DNA-PKcs on EJ outcomes, and also to disrupt DNA-PKcs with both M3814 treatment as well as genetic loss of DNA-PKcs (PRKDC-KO). For this, we used a MTAP/CDKN2B-AS1 deletion (MA-del) assay to examine various EJ outcomes via the chromosomal deletion rearrangement of the *CDKN2A* gene, a common deletion rearrangement found in various cancer types (Figure 2A)^46,47^. Expression of Cas9 and two sgRNAs induces two DSBs that target two loci that flank the *CDKN2A* gene (*MTAP* and *CDKN2B-AS1*). Subsequently, the deletion rearrangement is amplified using PCR, and amplicons are analyzed using deep sequencing. The reads are then aligned to the No Indel EJ product, i.e., repair of the distal DSB blunt ends without Insertions or Deletions. The analyzed reads are then categorized as No Indel EJ, Insertion, Deletion, and Complex Indel repair outcomes.

**Figure 2.**
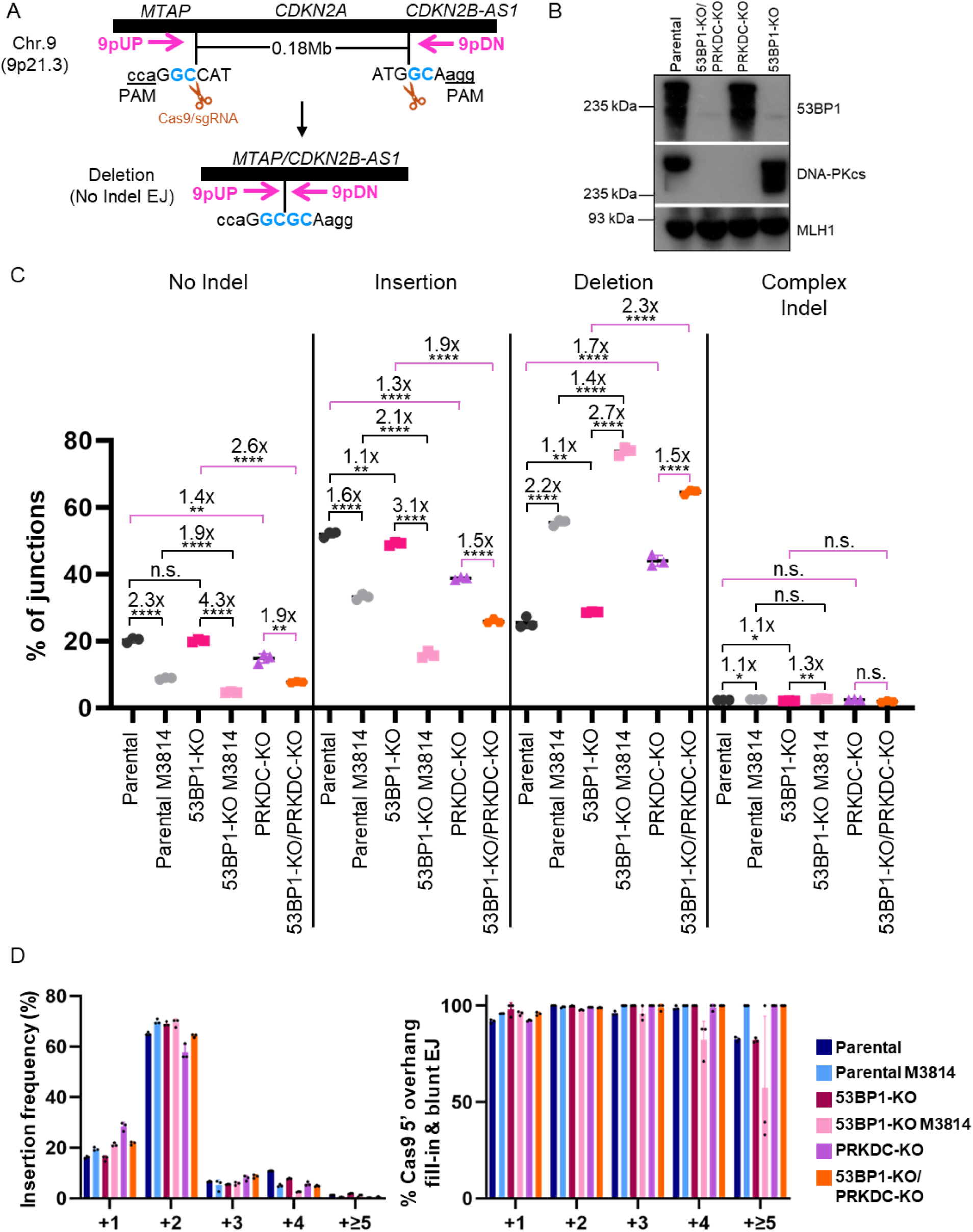
Combining loss of 53BP1 with either a DNA-PKcs inhibitor or genetic loss causes a decrease in DSB blunt EJ. **(A)** Schematic of the MTAP/CDK2NB1-AS1 deletion rearrangement reporter (MA-del), which induces two Cas9 DSBs that flank the CDKN2A locus. PCR is used to amplify the deletion rearrangement in preparation for deep sequencing analysis. (**B)** Immunoblot analysis of 53BP1 and DNA-PKcs in the Parental, single KO, and double KO cell lines. (**C)** Loss of 53BP1 alone has no obvious effect on indel types, but when combined with DNA-PKcs inhibition or loss, 53BP1 loss causes a decrease in No Indel EJ and Insertions, and in increase in Deletions. Shown are junction frequencies for No Indel EJ, Insertions, Deletions, and Complex Indels for each condition. n=3 biologically independent transfections. Statistics with unpaired t-test using Holm-Sidak correction. *P<0.05, **P>0.01, ***P>0.001, ****P>0.0001, n.s.=not significant. **(D)** Insertion events are likely associated with Cas9 5’ overhang fill-in and blunt DNA EJ in all conditions. Shown is the frequency of Insertion sizes, and percent Insertions consistent with Cas9 5’ overhang fill-in and blunt EJ for the experimental conditions shown in (B). n=3 biologically independent transfections.

We tested this assay on both Parental and 53BP1-KO cells with and without M3814 treatment, PRKDC-KO cells, as well as double knockout 53BP1-KO/PRKDC-KO cells (Figure 2B). Beginning with DNA-PKcs disruption, we found that M3814 treatment and DNA-PKcs loss each caused a decrease in No Indel EJ and Insertions, and an increase in Deletions (Figure 2C), which is consistent with a recent report^48^. In contrast, 53BP1 loss alone had no obvious effect on the relative frequencies of these EJ categories. However, in cells treated with M3814, loss of 53BP1 caused a decrease in No Indel EJ (1.9-fold) and Insertions (2.1-fold), and an increase in Deletions (1.4-fold) (Figure 2C). Similarly, with PRKDC-KO cells, loss of 53BP1 caused a decrease in No Indel EJ (1.9-fold) and Insertions (1.5-fold), and an increase in Deletions (1.5-fold) (i.e., comparing 53BP1-KO/PRKDC-KO vs. PRKDC-KO, Figure 2C). Finally, in all conditions, Complex Indels were rare (Figure 2C). These findings indicate that loss of 53BP1 alone does not cause an obvious effect on these EJ categories, but when combined with DNA-PKcs disruption via M3814 treatment or PRKDC-KO, causes a significant decrease in No Indel EJ and Insertion frequencies, and a significant increase in Deletion frequencies.

Because frequency patterns were similar between No Indel EJ and Insertion outcomes, we hypothesized that Insertion outcomes are caused by blunt DSB EJ. Specifically, we posited that Insertions mutations are caused by staggered Cas9 DSBs that cause 5’ overhangs that are filled in to generate blunt DSBs prior to EJ, as has been shown in several studies^10,48,49^. Insertions caused by this mechanism would result in specific sequences. Thus, we analyzed the Insertion sequences with the MA-del assay, and found that the vast majority of Insertions are consistent with this mechanism (Figure 2D, 2E), as found in a recent report^48^. For example, +2 nucleotide Insertions were the most frequent Insertion size and nearly all of the sequences consistent with staggered Cas9 DSBs that are filled in prior to blunt EJ, in all cellular conditions (Figure 2E). Thus, the Insertions appear to result from blunt EJ events, as are the No Indel EJ events. Accordingly, our findings indicate that 53BP1 is dispensable for blunt EJ unless combined with loss or inhibition of DNA-PKcs. We suggest that 53BP1 promotes DSB end synapsis during EJ, but this function is dispensable in the presence of DNA-PKcs.

### 53BP1 loss and DNA-PKcs disruption, alone and together, cause a similar shift in deletion patterns

Apart from blunt DSB EJ per se, we then assessed whether 53BP1 affects the pattern of deletion mutations, with or without disruption of DNA-PKcs. Namely, using the deletion mutations identified in the MA-del experiments above, we tested whether 53BP1 affects deletion size and/or microhomology use. Specifically, we analyzed the deletions to determine the frequency of deletions and the fraction of microhomology used for each deletion size (Figure 3A, B). From this analysis, we found that loss of 53BP1 alone caused a striking change in the Deletion pattern: a significant increase in −2, −7, −75 deletion sizes, which are associated with ≥2 nt microhomology, and a significant decrease in several deletion sizes without microhomology (i.e., 0, 1 nt.) (Figure 3A, B). There are a few notable exceptions to this pattern: loss of 53BP1 had no effect on the −13 deletion that is associated with microhomology, and interestingly caused a decrease in −17 and - 22 deletions associated with microhomology. We then compared the effects of 53BP1 loss to M3814 treatment, and found a very similar pattern. Namely, M3814 treatment caused a significant increase in −2, −7, −75 deletions with microhomology, and a decrease in several deletion sizes that do not utilize microhomology, similar to a recent report^48^ (Figure 3A, B). Also, as with 53BP1 loss, M3184 treatment had no effect on the −13 deletion associated with microhomology. However, for the −17 and −22 deletions associated with microhomology, M3814 treatment did not cause a decrease in these events, as was caused by 53BP1 loss. Notably, M3814 treatment in 53BP1-KO cells had very little effect on deletion size, with the exception of causing an increase in the −17 deletion. Finally, the −20 deletion associated with microhomology was only modestly affected by 53BP1 loss and M3814 treatment, and other deletion sizes with or without microhomology from –23 to −85 nt were relatively rare in each of these conditions (Figure 3A, B).

**Figure 3.**
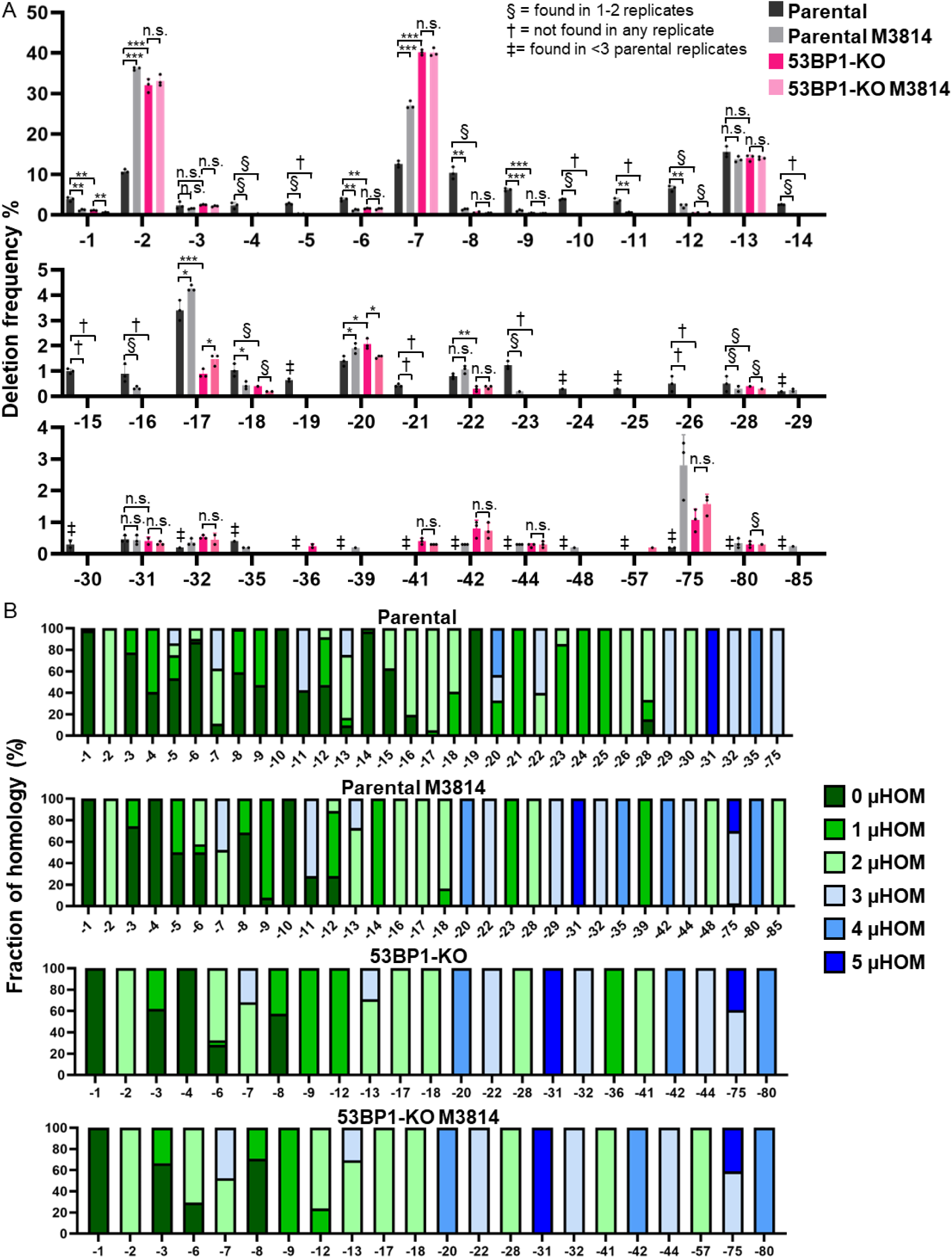
53BP1 loss and DNA-PKcs disruption, alone and together, cause a similar shift in deletion patterns. (**A)** Loss of 53BP1 causes an increase in −2 and −7 deletions, similar to inhibition of DNA-PKcs, and DNA-PKcs inhibition does not cause a further increase in 53BP1-KO cells. Shown are deletion sizes for the samples shown in Figure 2C. n=3 independent transfections. Statistics with unpaired t-test using Holm-Sidak correction. *P<0.05, **P<0.01, ***P<0.001, ****P<0.0001, n.s. = not significant. § = deletion size was only found in 1 or 2 replicates, † = deletion size was not found in any of the replicates, and ‡ = deletion size was found in <2 Parental replicates. (**B)** Microhomology use varies with distinct deletions. Shown is the fraction of microhomology used for each deletion size and experimental condition shown in (Figure 2C). n=3 biologically independent transfections.

We then evaluated genetic loss of DNA-PKcs. We found that such genetic loss (PRKDC-KO) caused a nearly identical effect on deletion mutation pattern as M3814 treatment, and combined loss of DNA-PKcs and 53BP1 (53BP1-KO/PRKDC-KO) was very similar to loss of DNA-PKcs alone (Supplementary Figure 2). Exceptions to this pattern were an increase in the −21 deletion without microhomology and a decrease in the −75 deletion with microhomology in the 53BP1-KO/PRKDC-KO vs. 53BP1-KO treated with M3814 (Figure 3A, B, Supplementary Figure 2).

### 53BP1 loss and DNA-PKcs disruption, alone and together, cause a similar increase in microhomology usage for deletions

The above analysis indicates that DNA-PKcs disruption and 53BP1 loss cause an increase in deletions with microhomology, and a reduction in deletions without microhomology. To examine microhomology another way, we compiled microhomology use for all deletions from the above MA-del experiment. From this analysis, 0-3 nt of microhomology were the most common in all conditions, but loss of 53BP1 and disruption of DNA-PKcs (M3814 and PRKDC-KO) caused a significant decrease in 0 and 1 nt microhomology deletions, and an increase in 2nt and 3 nt microhomology deletions (Figure 4). Furthermore, for these events, combined loss of 53BP1 with DNA-PKcs disruption was not different from the single knockouts. Deletions with 4 nt of microhomology were less common, but loss of 53BP1 and disruption of DNA-PKcs caused an increase in these events, except for PRKDC-KO/53BP1-KO cells. Altogether, these findings indicate that 53BP1 loss and DNA-PKcs disruption cause a similar increase in microhomology deletions, and that combined disruption of both factors is not additive.

**Figure 4.**
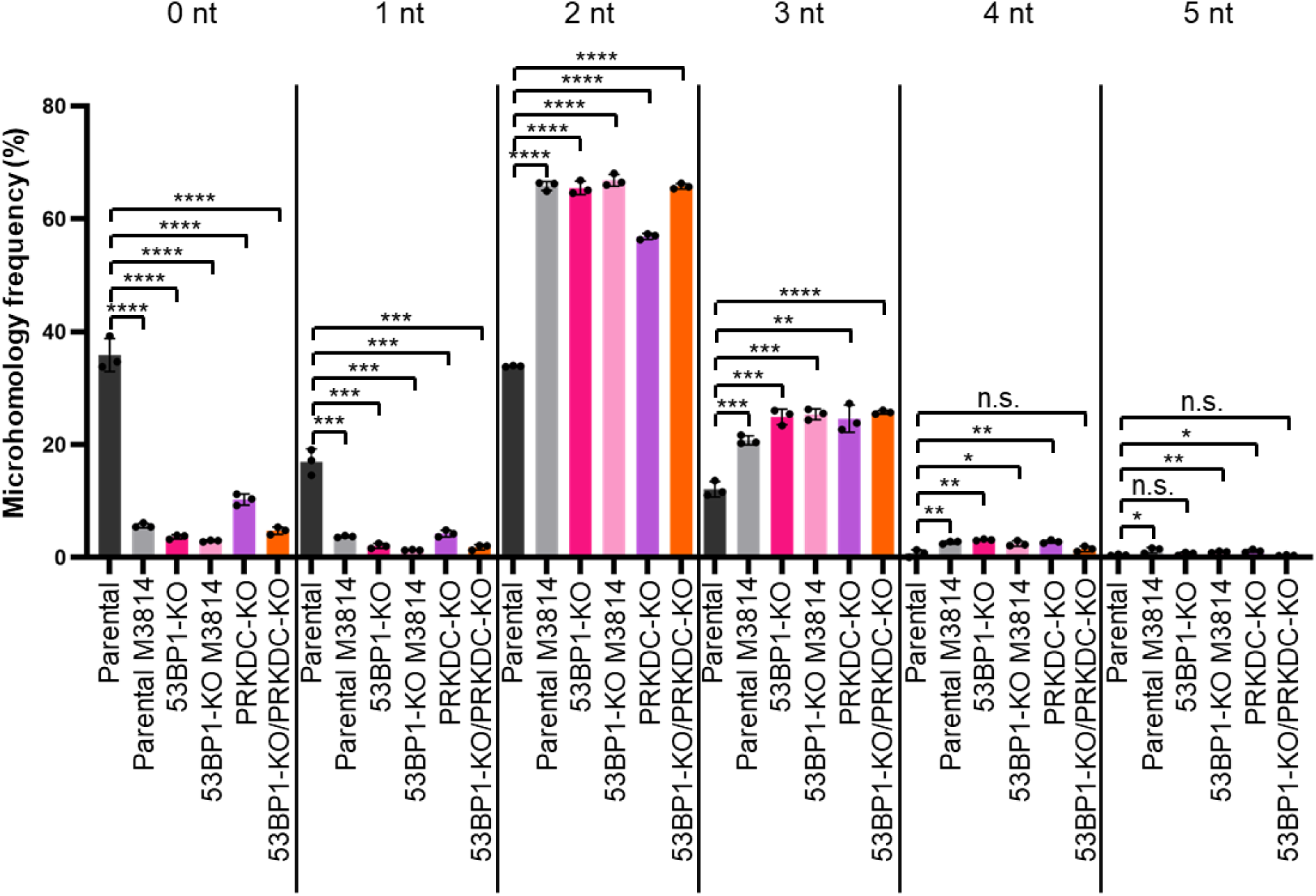
53BP1 loss and DNA-PKcs disruption, alone and together, cause a similar increase in microhomology usage for deletions. In all experimental conditions, the 2-nucleotide deletion is the most frequent deletion as compared to Parental. Shown is the frequency of microhomology used amongst all deletions for each experimental condition shown in (Figure 2C). n=3 biologically independent transfections. Statistics with unpaired t-test using Holm-Sidak correction. *P<0.05, **P<0.01, ***P<0.001, ****P<0.0001, n.s. = not significant.

### Loss of 53BP1 causes a modest decrease in DSB blunt EJ in XLF deficient cells

We then considered that loss of the C-NHEJ factor XLF might also affect the relative influence of 53BP1 on EJ outcomes, because both XLF and DNA-PKcs are implicated in DSB end synapsis^6,50,51^, and because loss of XLF revealed a role for 53BP1 for V(D)J recombination^23^. For this, we used the MA-Del assay to examine EJ outcomes in XLF-KO and double mutant 53BP1-KO/XLF-KO cells (Figure 5A, B). From this analysis, we found that XLF loss causes a reduction in No Indel EJ and Insertions (2.4-fold and 3.4-fold, respectively), and an increase in deletions (3.6-fold), which is consistent with its role in blunt DSB EJ from other studies^9,48^. We also found that the frequency of these EJ categories in the double mutant 53BP1-KO/XLF-KO was similar to the single XLF-KO (Figure 5B, No Indel EJ and deletions showed no difference, whereas Insertions were reduced 1.3-fold). These findings indicate that loss of XLF causes a reduction in No Indel EJ, which is not affected by 53BP1 loss.

**Figure 5.**
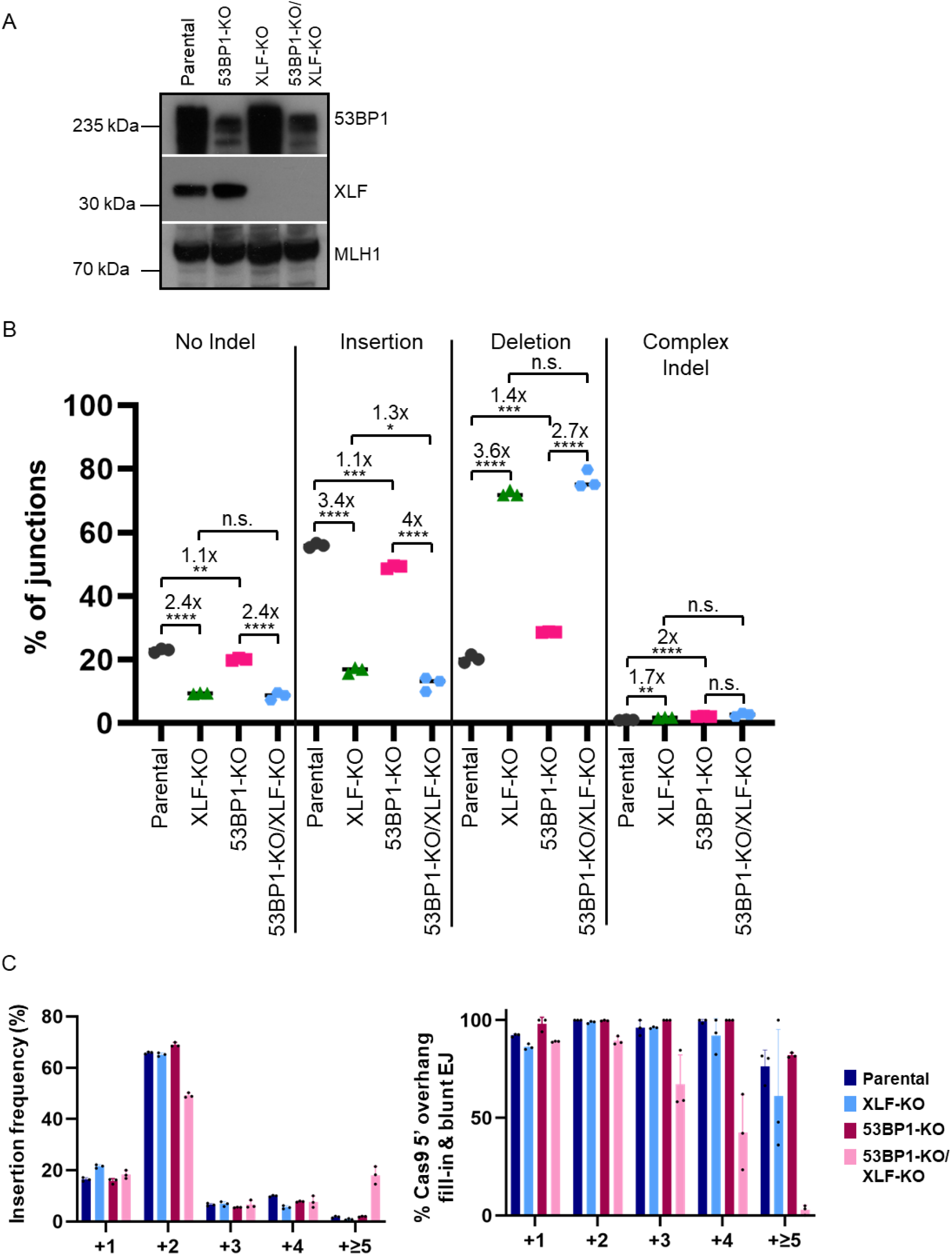
Loss of 53BP1 causes a modest decrease in DSB blunt EJ in XLF deficient cells. **(A)** Immunoblot analysis of 53BP1 and XLF in the Parental, 53BP1-KO and XLF-KO, and double KO cell lines. **(B)** Loss of XLF causes a reduction in No Indel EJ, which is not affected by 53BP1 loss, whereas combined loss of XLF and 53BP1 causes a modest reduction in Insertions, compared to XLF-KO. Shown are junction frequencies for No Indel EJ, Insertions, Deletions, and Complex Indels for each condition. n=3 biologically independent transfections. Statistics with unpaired t-test using Holm-Sidak correction. *P<0.05, **P>0.01, ***P>0.001, ****P>0.0001, n.s.=not significant. **(C)** Insertion events are likely associated with Cas9 5’ overhang fill-in and blunt DNA EJ are modestly reduced in 53BP1-KO/XLF-KO cells. Shown is the frequency of Insertion sizes, and percent Insertions consistent with Cas9 5’ overhang fill-in and blunt EJ for the experimental conditions shown in (B). n=3 biologically independent transfections

We then examined Insertions in more detail. Beginning with Insertion size, the general pattern of Insertion frequencies remains consistent between the various cell lines, however the 53BP1-KO/XLF-KO showed a 1.3-fold decrease in 2 nt Insertions, and a 10.4-fold increase in ≥5 nt Insertions, compared to the Parental cells (Figure 5C). Furthermore, the 53BP1-KO/XLF-KO showed a decrease in the frequency of Insertions consistent with staggered Cas9 cleavage, followed by 5’ overhang fill-in and blunt EJ: i.e., a 1.4-fold decrease in 3 nt Insertions, 2.3-fold decrease in 4 nt Insertions, and a 27.3-fold decrease in ≥5 nt Insertions (Figure 5C). Nonetheless, the larger Insertions in the 53BP1-KO/XLF-KO cells were relatively rare, i.e. most of the Insertions were similar to the Parental cell line. In summary, given that 53BP1-KO/XLF-KO showed a modest reduction in the frequency of Insertions (1.3-fold vs. XLF-KO), and a decrease in the insertions consistent with blunt DSB EJ, these results indicate that 53BP1 loss causes a modest decrease in blunt DSB EJ in XLF-KO cells.

### Loss of 53BP1 and XLF cause distinct deletion patterns, and the effect of 53BP1 dominates in the double mutant

We then assessed the effect of XLF loss with and without 53BP1 loss may affect the pattern of deletion mutations (i.e., deletion size and microhomology). From this analysis, we found that XLF loss caused a unique deletion pattern, similar to a recent study^48^. Specifically, loss of XLF did not have a substantial effect on the −2, −7, and −75 deletions associated with microhomology (Figure 6A, 6B), whereas as described above, loss of 53BP1 caused these events to increase (Figure 3A, 3B). In contrast, loss of XLF caused an increase in several microhomology associated deletions that range between −17 and −35 (Figure 6A, 6B). Furthermore, several deletion sizes without microhomology (i.e. 0-1 nt), were reduced by loss of XLF, which is similar to loss of 53BP1 (Figure 6A, B). These findings indicate that while XLF loss and 53BP1 loss each cause an increase in deletions associated with microhomology, the deletion sizes are different. Consistent with this notion, when we analyzed microhomology usage independent of deletion size, both XLF loss and 53BP1 loss caused an increase in microhomology usage (e.g. reduction in 0-1, and increase in 2-3 nt, Supplementary Figure 3).

**Figure 6.**
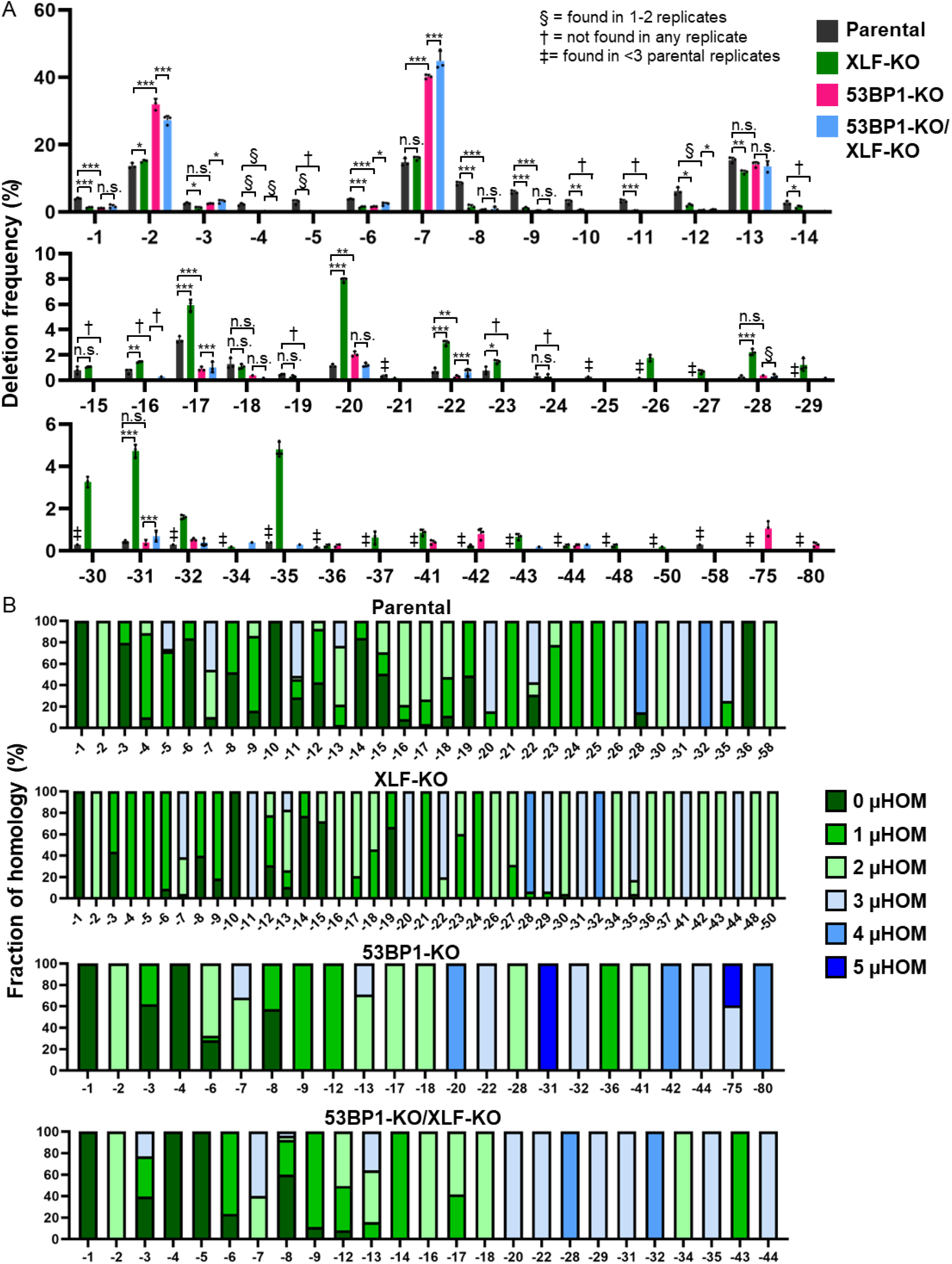
Loss of 53BP1 and XLF cause distinct deletion patterns, and the effect of 53BP1 dominates in the double mutant. **(A)** Combined loss of 53BP1 and XLF causes an increase in - 2 and −7 deletions, similar to loss of 53BP1 alone. Shown are deletion sizes for the samples shown in Figure 5B. n=3 independent transfections. Statistics with unpaired t-test using Holm-Sidak correction. *P<0.05, **P<0.01, ***P<0.001, ****P<0.0001, n.s. = not significant. § = deletion size was only found in 1 or 2 replicates, † = deletion size was not found in any of the replicates, and ‡ = deletion size was found in <2 Parental replicates. (**B)** Microhomology use varies with distinct deletions. Shown is the fraction of microhomology used for each deletion size and experimental condition shown in (Figure 5B). n=3 biologically independent transfections.

Interestingly, the deletion pattern for cells with combined loss of 53BP1 and XLF largely mimics the deletion pattern of loss of 53BP1 alone. Namely, compared to Parental cells, combined loss of 53BP1 and XLF caused a significant increase in −2 and −7 deletions associated with microhomology, similar to 53BP1-KO (Figure 6A, 6B). Furthermore, the deletions associated with microhomology ranging between −17 and −35 that were substantially elevated in XLF-KO, were similar between 53BP1-KO and 53BP1-KO/XLF-KO. As an exception to this pattern, combined loss of 53BP1 and XLF failed to cause an increase in the −75 deletion size, which was similar to XLF-KO, not 53BP1-KO. Altogether, these findings indicate that loss of 53BP1 and XLF cause distinct deletion frequency patterns to each other, and the pattern caused by 53BP1 loss is largely dominant over XLF loss in the double knockout.

### Combined loss of RIF1 and DNA-PKcs inhibition causes a decrease in No Indel EJ

A key downstream effector protein of 53BP1 is RIF1^27–38^. Thus, we considered that RIF1 and 53BP1 may have similar effects on EJ outcomes. For this, we generated a HEK293 RIF1-KO cell line with the EJ7-GFP reporter for No Indel EJ (Figure 7A). With this reporter, we found that loss or RIF1 did not obviously affect the frequency of No Indel EJ, although transient expression of RIF1 in the RIF1-KO caused a modest (1.4-fold) increase (Figure 7A). In contrast, in cells treated with the M3814 DNA-PKcs inhibitor, loss of RIF1 caused a substantial (3.3 fold) decrease in No Indel EJ, which was recovered with transient expression of RIF1-WT (Figure 7A). As a control, we found that M3814 treatment inhibits DNA-PKcs kinase activity similarly in RIF-KO and Parental cells (Supplementary Figure 1). These findings indicate that RIF1 promotes No Indel EJ under conditions of DNA-PKcs inhibition, which is similar to the above findings with 53BP1.

**Figure 7.**
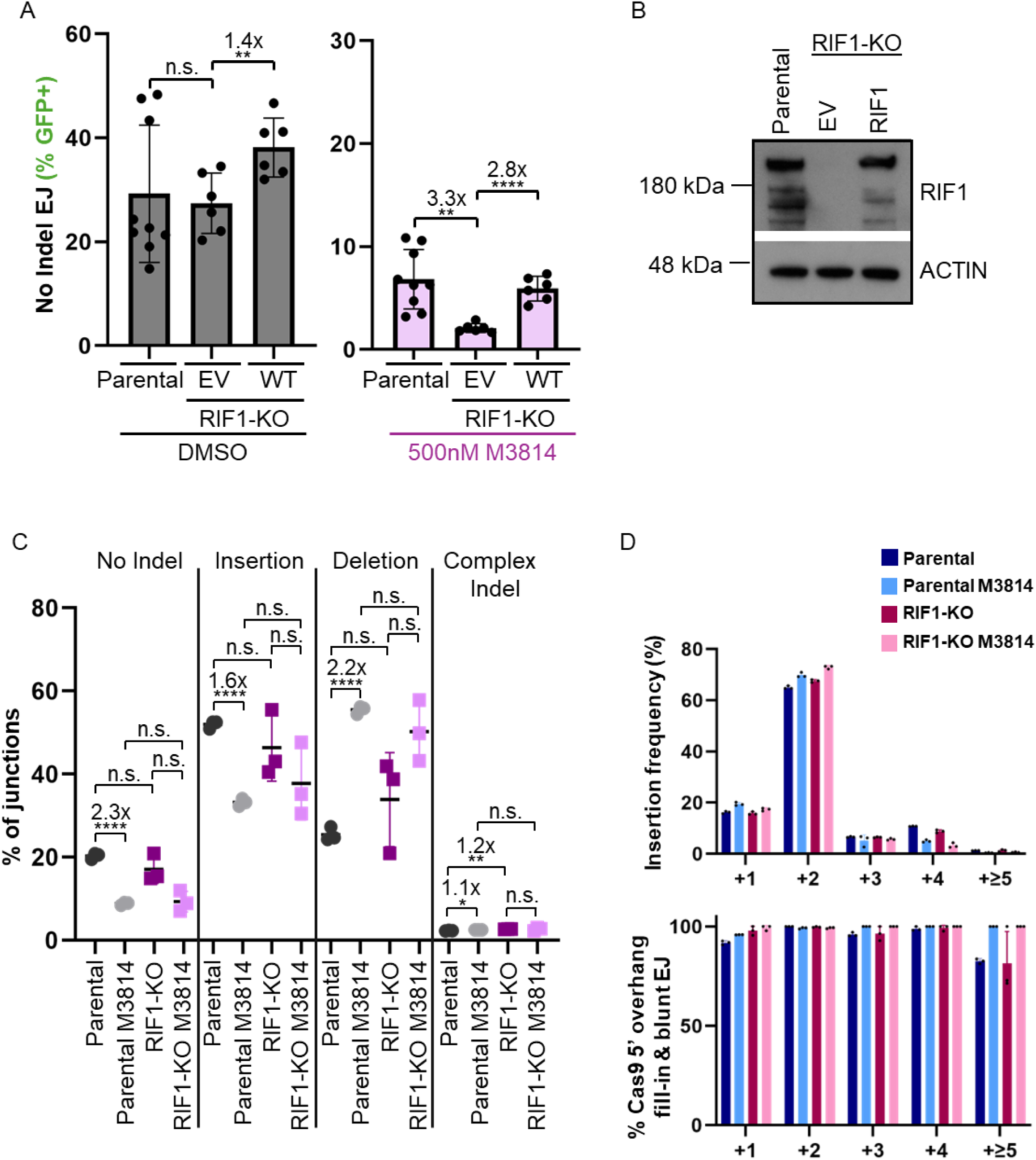
Combined loss of RIF1 and DNA-PKcs inhibition causes a decrease in No Indel EJ. **(A)** Loss of RIF1 alone is not important for No Indel EJ, but becomes important with inhibition of DNA-PKcs. n=6, except Parental n=9. Statistics with unpaired t-test using Holm-Sidak correction. **P<0.01, ****P<0.0001, n.s. = not significant. Parental values are those seen in Figure 1B. **(B)** Immunoblot analysis of RIF1 for the cell lines and conditions shown in (B). **(C)** Loss of RIF1 alone has no obvious effect on EJ junction patterns. Shown are junction frequencies for No Indel EJ, Insertions, Deletions, and Complex Indels for each condition. n=3 biologically independent transfections. Statistics with unpaired t-test using Holm-Sidak correction. *P<0.05, **P>0.01, ****P>0.0001, n.s. = not significant. Parental values are those seen in Figure 2C. **(D)** Insertion events are likely associated with Cas9 5’ overhang fill-in and blunt DNA EJ in all conditions. Shown is the frequency of Insertion sizes, and percent Insertions consistent with Cas9 5’ overhang fill-in and blunt EJ for the experimental conditions shown in (C). n=3 biologically independent transfections.

We then used the MA-del assay to examine the effect of loss of RIF1 and DNA-PKcs inhibition on various EJ outcomes. We found that loss of RIF1, alone, and combined with DNA-PKcs inhibition had no significant effect on the categories of EJ junctions (i.e., No Indel, Insertion, or Deletion outcomes, Figure 7C). Additionally, we also found that loss of RIF1 had no obvious effect on the pattern of Insertions (i.e., length and % consistency with Cas9 staggered DSBs, Figure 7D). Thus, while loss of RIF1 combined with DNA-PKcs inhibition caused a reduction in No Indel EJ with the EJ7-GFP reporter, we did not observe this difference with the MA-del assay. We speculate that the MA-del assay may be less sensitive to detect defects in No Indel EJ vs. EJ7-GFP. Consistent with this notion, XLF loss has been shown to cause a greater defect on No Indel EJ with the EJ7-GFP reporter vs. the MA-del assay^48^.

### RIF1 loss causes a similar shift in deletion patterns as DNA-PKcs disruption

Even though RIF1 loss did not affect the frequency of EJ categories with the MA-del assay, we considered that it may affect the deletion pattern (i.e., size and/or microhomology use), as we found with 53BP1. From analyzing the deletions in the above experiment, we found that loss of RIF1 causes an increase in microhomology-associated −2, −7, and −75 deletions (Figure 8A, 8B). Furthermore, M3814-treatment of RIF1-KO cells caused a slight increase in −2 deletion size frequency, but not for the −7 or −75 deletions. (Figure 8A). This pattern is very similar to that of loss of 53BP1, which itself is similar to disruption of DNA-PKcs (Figure 3, Supplementary Figure 2). Finally, from analysis of microhomology use from deletions, we found that loss of RIF1 causes an increase in microhomology that is similar to M3814 treatment, and combined RIF1 loss with M3814 treatment is similar to RIF1 loss alone (Supplementary Figure 4). These findings indicate that RIF1 has a similar role to limit microhomology-associated deletions as 53BP1 and DNA-PKcs.

**Figure 8.**
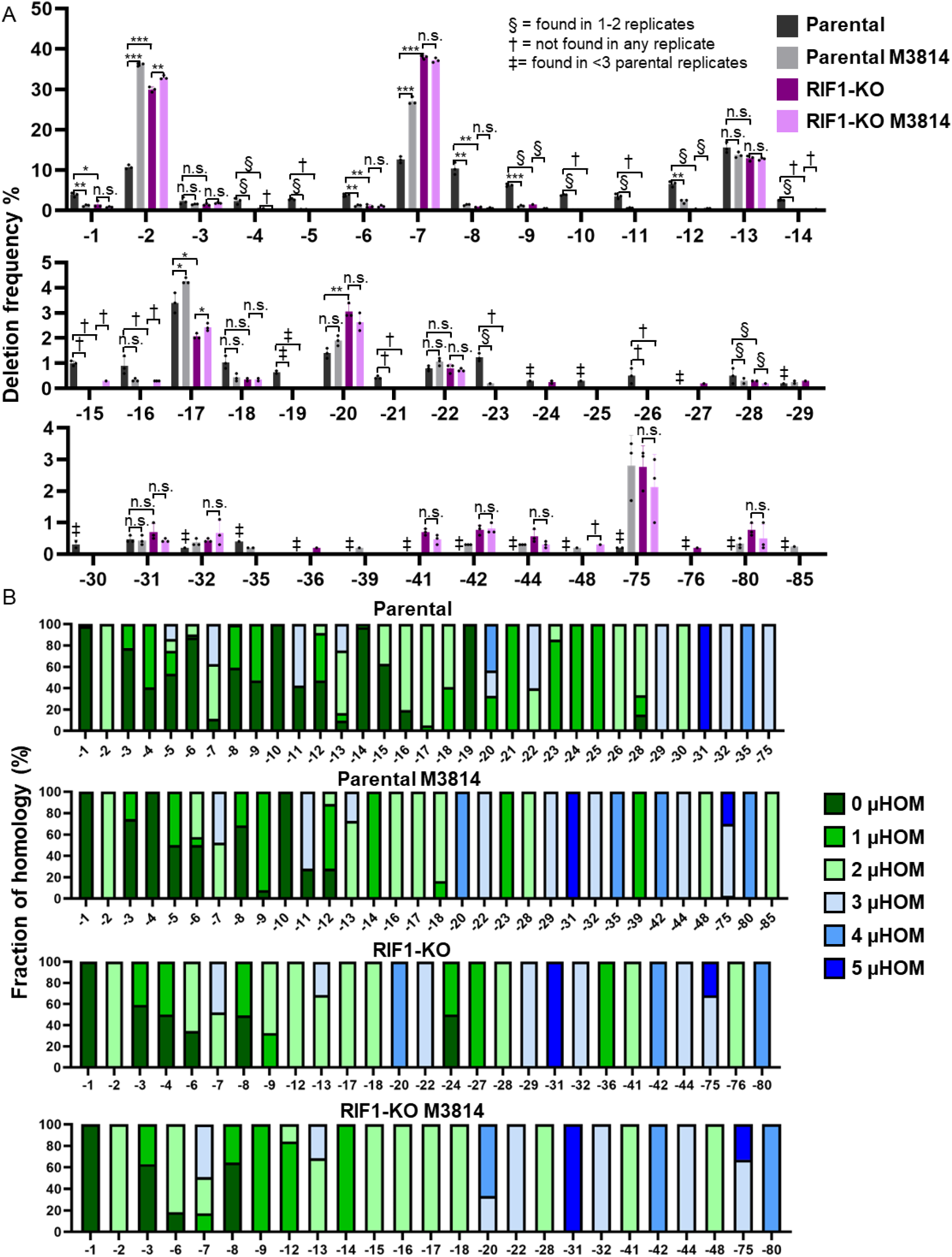
RIF1 loss causes a similar shift in deletion patterns as DNA-PKcs disruption. **(A)** Loss of RIF1 causes an increase in −2 and −7 deletions, similar to inhibition of DNA-PKcs. Shown are deletion sizes for the samples shown in (Figure 5C). n=3 independent transfections. Statistics with unpaired t-test using Holm-Sidak correction. *P<0.05, **P<0.01, ***P<0.001, ****P<0.0001, n.s. = not significant. Parental values are those seen in (Figure 3A). § = deletion size was only found in 1 or 2 replicates, † = deletion size was not found in any of the replicates, and ‡ = deletion size was found in <2 Parental replicates. (**B)** Microhomology use varies with distinct deletions. Shown is the fraction of homology used for each deletion size and experimental condition shown in (Figure 5C). N=3 biologically independent transfections. Parental values are those seen in (Figure 3B).

As a control for all MA-del assay experiments, we also examined the total frequency of the deletion rearrangement, using a quantitative PCR (qPCR) assay. Namely, we performed qPCR of the MA-del deletion rearrangement, normalized to a control PCR in the *MTAP* locus, each quantified using a fluorescent probe. As expected, we found that the qPCR signal for the MA-deletion rearrangement was dependent on transfection of the Cas9/sgRNAs that target the *MTAP* and *CDKN2B-AS1* loci (>80-fold increase, Supplemental Figure 2). Notably, none of the M3814 treatments and genetic disruptions caused a significant decrease in deletion frequency (Supplemental Figure 2). Although, a few treatments and genetic disruptions caused an increase in deletion frequency, compared to the Parental control (Supplemental Figure 2). In summary, the deletion frequency was readily detected, and not obviously reduced, with all the variants of genetic disruptions and M3814 treatments.

### Influence of 53BP1, RIF1, and DNA-PKcs on HDR and radiosensitivity

Finally, we examined other aspects of DSB repair: gene editing via HDR and radiosensitivity. Inhibition of DNA-PKcs has been shown to cause elevated HDR and cause radiosensitization, which have clinical applications^42,44,52,53^. Thus, we sought to define how 53BP1 and RIF1 affect the relative influence of DNA-PKcs disruption on these processes.

To examine HDR, we used the LMNA-HDR assay, which involves inducing a Cas9 DSB at the *LMNA* gene along with a donor template plasmid, which if used for HDR causes an *LMNA-mRuby2* fusion gene, encoding a fluorescent protein that can be measured by flow cytometry (Figure 9A). Using this assay, we found that M3814 treatment alone caused a substantial increase in HDR (2.8-fold), whereas genetic loss of DNA-PKcs caused a mild increase (1.3-fold), similar to prior studies (Figure 9B). Loss of 53BP1 caused an increase in HDR alone (4.1-fold), in M3814 treated cells (2.2-fold), and in with DNA-PKcs loss (1.6-fold, i.e. 53BP1-KO/PRKDC-KO vs. PRKDC-KO, Figure 9B). Furthermore, transient expression of 53BP1 caused a decrease in HDR in each of these scenarios. Thus, 53BP1 inhibits HDR in both DNA-PKcs proficient and deficient cells, although in the latter, the fold effect of 53BP1 loss on HDR is diminished (Figure 9B).

**Figure 9.**
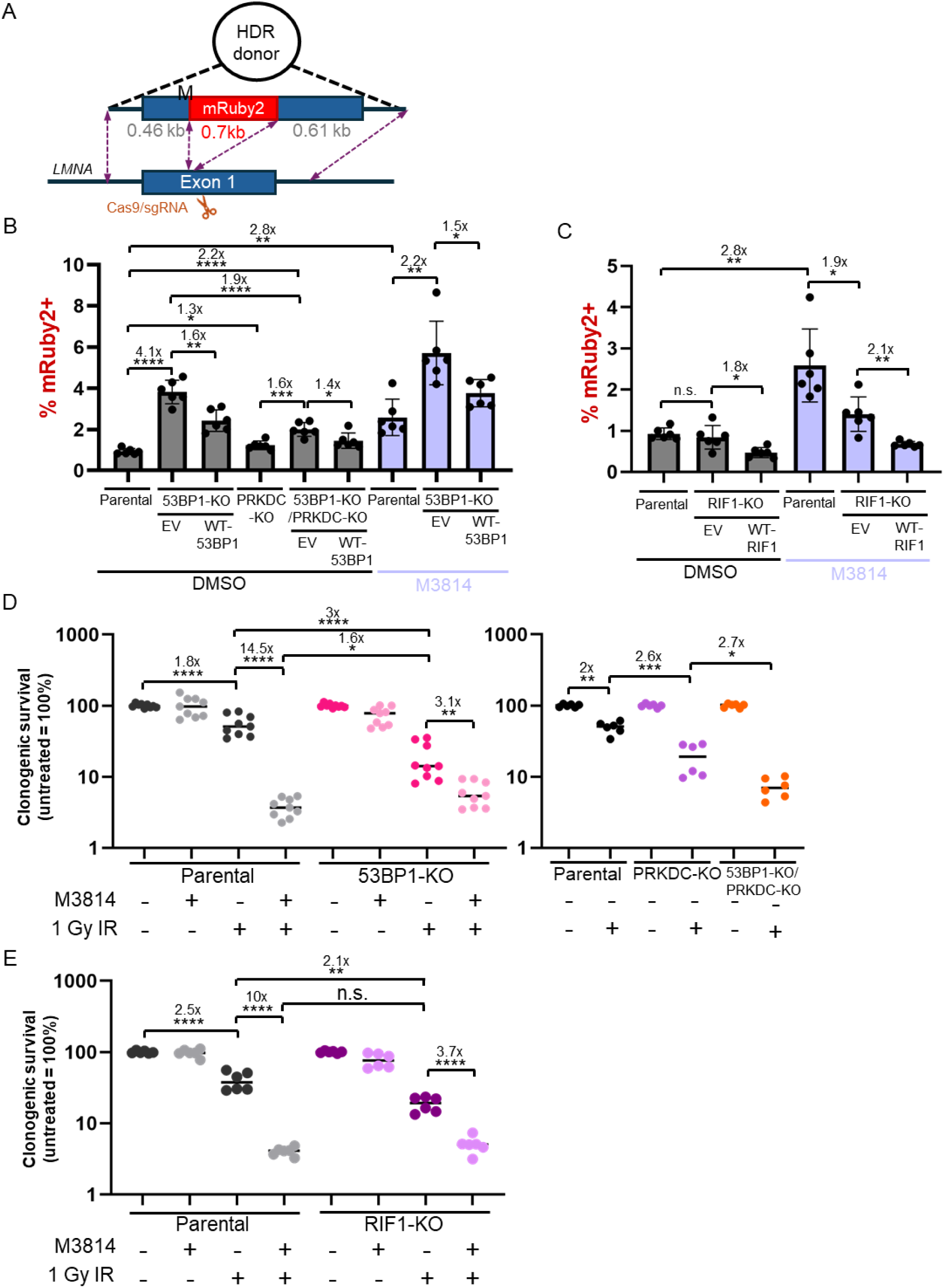
Influence of 53BP1, RIF1, and DNA-PKcs on HDR and radiosensitivity. **(A)** Schematic of the LMNA-HDR reporter that involves a Cas9/sgRNA expression vector that induces a single DSB in *LMNA* exon 1, as well as plasmid donor, where HDR yields expression of mRuby2 from the *LMNA* locus. mRuby2 frequencies are normalized to transfection efficiency with parallel GFP transfections. (**B)** 53BP1 loss causes an increase in HDR in both DNA-PKcs proficient and deficient cells. n=6 biologically independent transfections. Statistics with unpaired t-test using Holm-Sidak correction. *P<0.5, **P<0.01, ****P<0.0001. (**C)** RIF1 loss does not obviously increase HDR, but RIF1 expression inhibits HDR independent of DNA-PKcs kinase inhibition. n=6 biologically independent transfections. Statistics with unpaired t-test using Holm-Sidak correction. *P<0.5, **P<0.01, n.s.=not significant. **(D)** 53BP1 loss causes radiosensitivity that less than DNA-PKcs kinase inhibition, but similar to DNA-PKcs loss. Cells were treated with M3814 or vehicle control (DMSO), and 0 Gy or 1 Gy IR dose, and plated to form colonies. Fraction clonogenic survival was determined relative to the untreated control (DMSO 0 Gy) for each cells line. n=6. Statistics with unpaired t-test using Holm-Sidak correction. *P<0.05, **P>0.01, ***P>0.001, ****P>0.0001. **(E)** Loss of RIF1 causes radiosensitivity that is less than DNA-PKcs kinase inhibition. Cells were treated and analyzed as in D. n=6. **P>0.01, ****P>0.0001, n.s.=not significant.

The effects of RIF1 on the LMNA-HDR assay were more complicated. Loss of RIF1 alone had no significant effect on HDR, and when combined with M3814 treatment caused a modest decrease (Figure 9C). However, transient expression of RIF1 caused a decrease in HDR both with and without M3814. We conclude that RIF1 loss does not cause an obvious increase in HDR by this assay, although expression of RIF1 appears to cause a decrease.

Lastly, we examined clonogenic survival (colony formation) following ionizing radiation (IR). We selected one dose of IR (1 Gy), based on prior studies showing significant radiosensitization via M3814 in HEK293 cells at this dose. Indeed, M3814 treatment of Parental cells caused a marked hypersensitivity to IR (14.5-fold, Figure 9D). Loss of 53BP1 alone caused a 3-fold hypersensitivity to IR, compared to Parental cells, and treating 53BP1-KO cells with M3814 caused a further 3.1-fold hypersensitivity to IR (Figure 9D). Thus, while M3814 treatment caused hypersensitivity to 53BP1-KO cells, the fold effect was less as compared to Parental cells. Indeed, 53BP1-KO cells treated with M3814 were only modestly radiosensitive compared to M3814-treated Parental cells (1.6-fold, Figure 9D). We then tested combined genetic loss of 53BP1 and DNA-PKcs (PRKDC-KO), finding that PRKDC-KO cells show a 2.6-fold hypersensitivity to IR vs. Parental cells, and the double mutant (53BP1-KO /PRKDC-KO) was a further 2.7-fold hypersensitive compared to the PRKDC-KO single mutant (Figure 9D). Loss of RIF1 alone caused a 2.1-fold hypersensitivity to IR, and M3814 treatment of RIF1-KO cells caused a further 3.7-fold hypersensitivity (Figure 9E). In summary, loss of 53BP1 and RIF1 cause IR sensitivity, and while addition of M3814 treatment causes further radiosensitivity in these cells, the effects are not additive. In contrast, combining genetic loss of 53BP1 and DNA-PKcs causes approximately additive hypersensitivity to IR.

## DISCUSSION

53BP1 is a key DNA damage response factor recruited to DSBs through interactions with chromatin marks that flank DSBs^27^. 53BP1 has been shown to affect DSB repair by inhibiting HDR, particularly in BRCA1-deficient cells, mediating class switch recombination during antibody maturation, as well as promoting fusion of deprotected telomeres^21–27^. However, the influence of this factor on affecting diverse EJ outcomes, and its functions in DSB repair in relation to C-NHEJ, have been unclear. We have focused on the genetic relationship between 53BP1 and DNA-PKcs, because both factors have been implicated in DSB end synapsis and regulation of DSB end processing^3–7,21–27,40,54^. Furthermore, defining the interplay with DNA-PKcs is significant, as DNA-PKcs kinase inhibitors are in clinical development^42,44,52,53^. We also examined RIF1, which is a key effector of 53BP1^27–38^.

From analysis of blunt DSB EJ, which is dependent on several C-NHEJ factors (e.g., XRCC4), we found that 53BP1 is dispensable but plays a backup role in cells disrupted for DNA-PKcs (both kinase inhibition and genetic loss, Fig 10). This decrease in blunt DSB EJ is associated with a shift toward deletion mutations. Thus, 53BP1 appears to promote blunt DSB EJ and suppress EJ involving end processing, but these roles are masked by the function of DNA-PKcs. DNA-PKcs promotes DSB end synapsis in a long-range complex via interactions with the DNA-PKcs dimer and interactions with Ku80 on the opposing DSB end^5–8^. Subsequently this long-range complex can transition to a short-range complex without DNA-PKcs^5–8^. In this short-range complex, DSB ends are positioned for ligation^5–8^. We suggest that 53BP1 can also assist with end synapsis to stabilize the long-range complex and/or short-range complex, but serves as a backup to synapsis mediated by DNA-PKcs. As a possible mechanism for mediating end synapsis, 53BP1 can oligomerize and form large condensates^41^, as well as form microdomains that are important for chromosome condensation at DSBs^40^. Alternatively, 53BP1 could mediate direct interactions with the C-NHEJ complex to support end synapsis.

**Figure 10.**
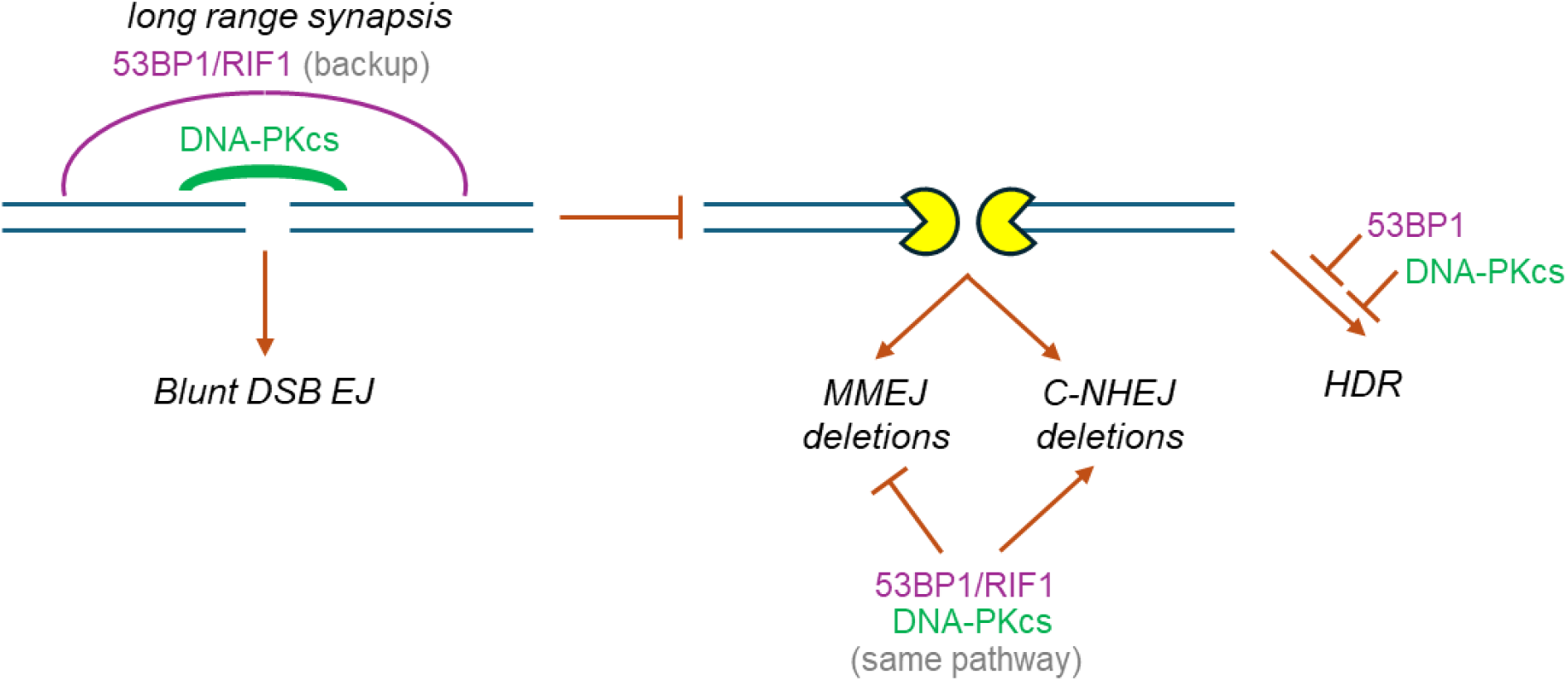
Model of genetic relationship between 53BP1/RIF1 and DNA-PKcs on diverse DSB repair outcomes.

While 53BP1 loss did not affect the frequency of deletion mutations, it had a substantial effect on the type of deletions. Specifically, loss of 53BP1 caused an increase in deletions with microhomology, and conversely a decrease in those without microhomology (Fig 10). In this case, the effect of 53BP1 loss was similar to that of DNA-PKcs disruption (both genetic loss and kinase inhibition), and the combined disruption did not have an additive effect. Thus, 53BP1 and DNA-PKcs appear to function in the same pathway to affect the type of deletions. Our findings are consistent with a recent report that 53BP1 loss (in *Ku70-/-* mouse cells) causes an increase in microhomology usage during V-J recombination^55^. We speculate that 53BP1 and DNA-PKcs could be important to promote end processing events that are tightly linked to completion of repair by C-NHEJ. This model is supported by studies of the kinetics of C-NHEJ repair of IR-induced damage in G1, which appears to occur in a fast phase and a slow phase, the latter of which may involve DSB end processing^56^. Notably, while XLF loss also caused an increase in microhomology deletions, the deletion sizes were distinct from those affected by DNA-PKcs and 53BP1^48^. Furthermore, with combined loss of 53BP1 and XLF, the deletion pattern caused by loss of 53BP1 appears to dominate. The mechanisms that drive these distinct microhomology deletion patterns are unclear. We suggest that further defining the factors that influence the type of deletion mutation during EJ will provide insight into such mechanisms.

The effects of disrupting 53BP1 on EJ were similar for loss of RIF1, which is a direct effector protein that binds phosphorylated 53BP1^27–38^. Namely, loss of RIF1 caused a reduction in No Indel EJ when combined with DNA-PKcs kinase inhibition and affected the type of deletion mutation similarly to 53BP1 loss. Although, the effect on No Indel EJ was somewhat more modest for RIF1, in that the effect was detectable with the EJ7-GFP assay, but not the junction patterns of the MA-del assay. We suggest that RIF1, like 53BP1, mediates end synapsis as a backup to DNA-PKcs (Fig 10). Consistent with a role in end synapsis, RIF1 appears to stabilize condensed chromatin at DSBs, and promote 53BP1 macrodomains that may be involved in such stabilization^40^. While RIF1 loss did not obviously affect the categories of EJ outcomes the MA-del assay (i.e., No Indel EJ, Deletions, Insertions, and Complex Indels), loss of RIF1 indeed yielded a shift to microhomology deletions that is similar to loss of 53BP1 and disruption of DNA-PKcs. Thus, as with 53BP1, RIF1 appears to suppress microhomology EJ deletions, which is consistent with the notion that RIF1 is a key effector protein of 53BP1^27–38^. As mentioned above, defining additional factors that affect the type of deletion mutation will provide insight into the mechanism of 53BP1/RIF1-mediated suppression of microhomology EJ deletions.

We also examined the influence of 53BP1, RIF1, and DNA-PKcs on HDR. Loss of 53BP1 caused an increase in HDR in both DNA-PKcs proficient and deficient cells. Conversely, DNA-PKcs kinase inhibition also caused an increase in HDR in both 53BP1 proficient and deficient cells. However, the fold-effects of 53BP1 loss and DNA-PKcs kinase inhibition were less than additive, which might indicate a partial overlap in function. For example, loss of 53BP1 and DNA-PKcs kinase inhibition might cause an increase in HDR via distinct mechanisms, yet the combination could cause a maximal possible increase in HDR. Interestingly, DNA-PKcs kinase inhibition caused a much greater increase in HDR vs. genetic loss, which reflects that kinase inhibition does not always have the same effect as loss. Indeed, DNA-PKcs kinase inhibition causes a more severe mouse phenotype vs. the genetic knockout, likely because DNA-PKcs kinase inhibition stabilizes its interaction with DNA ends^57^. Indeed, these findings with HDR are consistent with reports that DNA-PKcs bound to DNA ends are a signal for initiation of DSB end resection via the MRE11 complex and CtIP^54^.

Finally, we examined the influence of disrupting 53BP1, RIF1, and DNA-PKcs on radiosensitivity. We found that DNA-PKcs kinase inhibition caused marked radiosensitivity, which was greater than the radiosensitivity caused by genetic loss of DNA-PKcs. Furthermore, the effects of genetic loss of DNA-PKcs, 53BP1, and RIF1 were similar to each other. Notably, we also found that DNA-PKcs kinase inhibition had the greatest overall effect on DSB repair outcomes: causing an increase in HDR, a decrease in blunt DSB EJ, and an increase in microhomology deletions. We speculate that the combination of these diverse effects on DSB repair outcomes contribute to the substantial radiosensitivity caused by DNA-PKcs kinase inhibition. Namely, these defects in EJ could cause toxic mutations and chromosomal rearrangements. Furthermore, elevated HDR could cause repair before sister chromatid synthesis, resulting in use of ectopic donors that can also cause chromosomal rearrangements. In contrast, genetic loss of DNA-PKcs, 53BP1, and RIF1, each had lesser effects on DSB repair outcomes vs. DNA-PKcs kinase inhibition. For example, loss of 53BP1 alone failed to cause a defect in blunt DSB EJ, but caused similar effects as DNA-PKcs kinase inhibition on causing an increase in microhomology deletions and HDR. In conclusion, we suggest that 53BP1/RIF1 and DNA-PKcs show distinct genetic interactions depending on specific DSB repair outcomes, which together are important for radioresistance.

## METHODS

### Cell lines and plasmids

For sgRNA and Cas9 expression, the px330 plasmid was used (deposited by Dr. Feng Zhang, Addgene 42230)^58^. All sequences for the sgRNAs used in this study, including the sgRNAs for the EJ7-GFP assay (7a and 7b), and the MA-del assay (MTAP and CDKN2B-AS1) are found in Supplemental Table 1. The pCAGGS-53BP1 expression vector was described previously^45^, and the pCAGGS-RIF1 expression vector used the RIF1 coding sequence from pDEST pcDNA5-FRT/TO-eGFP-RIF1 (deposited by Dr. Daniel Durocher, Addgene 52506)^29^, and the empty vector control is pCAGGS-BSKX^59^. Plasmids for the LMNA-HDR assay (LMNA Cas9/sgRNA and LMNA-mRuby2-Donor plasmids) were provided by Dr. Jean-Yves Masson, and were previously described^60^.

The HEK293 EJ7-GFP cell line, the XLF-KO and PRKDC-KO were previously described^42,61^. 53BP1-KO, RIF1-KO, 53BP1-KO/XLF-KO, and 53BP1-KO/PRKDC-KO cell lines were generated using Cas9/sgRNAs targeting 53BP1 and RIF1, each using one sgRNA for RIF1, and two sgRNAs for 53BP1. To generate the knockout cell lines, cells were co-transfected with the Cas9/sgRNA and pgk-puro plasmid, transfected cells were enriched using puromycin treatment, and were then plated at low density to isolate and screen individual clones for genetic disruption via immunoblotting.

### DSB Reporter Assays

To test the EJ7-GFP reporter assay, HEK293 cells were seeded in a 24-well plate coated with poly-lysine at 0.5×10^5^ cells/well. The next day, cells were transfected with 200 ng of sgRNA/Cas9 plasmids (7a and 7b), and 50 ng of 53BP1, RIF1, or EV control plasmid. To test transfection efficiency, parallel transfections were performed with 200 ng of GFP expressing plasmid (pCAGGS-NZE-GFP) and 200 ng of EV with respective amounts of 53BP1, RIF1, and EV. For each well, all transfections used 1.8 μL of Lipofectamine 2000 (Thermofisher) and 0.5 mL of antibiotic-free media. To test the LMNA-HDR assay, LMNA Cas9/sgRNA and LMNA-mRuby2-Donor plasmids were used in place of the two sgRNA/Cas9 plasmids (200 ng each). Cells were incubated for 4 hours with the transfection agents, washed, and treated with media containing 500 nM M3814 (i.e., Nedisertib, Selleckchem #S8586) or vehicle (Dimethyl Sulfoxide, DMSO). All wells had the same total amount of DMSO in each experiment. 3 days after the transfection, cells were analyzed using flow cytometry (ACEA Quanteon, Agilent NovoExpress Version 1.5.0), as described^42^.

For the MA-del assay, cells were seeded in a 6-well plate coated with poly-lysine. Cells were transfected with 800 ng each of MTAP and CDK2NB-AS1 Cas9/sgRNA plasmids, as well as 400 ng of pgk-puro plasmid. Each well used 7.2 μL of Lipofectamine 2000 and 2 mL of antibiotic-free media. As in the EJ7 assay, cells were incubated for 4 hours with transfection agents, washed, and treated with media containing M3814 or DMSO. The day after transfection, cells were plated into M3814- or DMSO-containing complete media with 5 μg/mL puromycin for 2 days to enrich for transfected cells via puromycin selection and then expanded into media without puromycin for 3 days. Genomic DNA isolation was conducted as described^59^. PCR amplification (Platinum HiFi Supermix, Thermo Fisher) of the MTAP-CDK2NB1 rearrangement used the MAfusion1UP and MAfusion1DN primers.

The amplicons were subjected to deep sequencing using the Amplicon-EZ service (Azenta) and their SNP/INDEL detection pipeline, which uses the Burrows-Wheeler Aligner (BWA) to align the reads to the predicted No Indel EJ junction sequence based on the sgRNA/Cas9 cut sites. The indel categories were identified subsequent to this alignment step: WT (No Indel EJ product), base changes, deletions (continuous or discontinuous, and with or without base changes), Insertions (inserted nucleotides without deletions, and with or without base changes), and Insertion and deletion (both Insertion and deletion, and with or without base changes). From this analysis, EJ outcomes were categorized as No Indel EJ, Insertions, Deletions, or Complex Indels. Then, the total reads in each category were used to assess their frequency. Insertion and deletion sequences were analyzed individually by manual alignment, for all read sequences representing at least 0.1% of the combination of Insertions and complex indels, and at least 0.1% of the deletion reads, respectively. Some deletion events showed nucleotides consistent with staggered Cas9 cleavage, which were used to assign breakpoints for deletion size and microhomology. For each condition, amplicon sequencing was performed on 3 independent transfections, and frequencies represent the mean ± SD.

To quantify the deletion rearrangement by qPCR, two sets of reactions were performed, both using the probe primer (MAfusionHYB), and with MAfusion1UP and MAfusion1DN for the deletion rearrangement, and MAfusion1UP and MTAPctrlDN1 for the control reaction. The qPCR was performed with iTaq Universal Probes Supermix and the CRX Connect Real-Time PCR Detection System (BioRad). The relative levels of the deletion rearrangement were determined using the cycle threshold (Ct) value for the deletion rearrangement reaction and then subtracted by the average Ct value for the control reaction (ΔCt). This value was then normalized to the ΔCt value from Parental cells analyzed in parallel, to calculate the 2−ΔΔCt value.

### Immunoblotting

To conduct immunoblot analysis, cells were seeded in a poly-lysine coated 6-well plate and were transfected as previously described in the reporter assays, with the exception that EV plasmid (pCAGGS-BSKX) was used instead of the sgRNA/Cas9 plasmid(s). Transfected cells were scraped from each well, lysed with ELB (250 mM NaCl, 5 mM EDTA, 50 mM Hepes, 0.1% (v/v) Ipegal, and Roche protease inhibitor), and then were sonicated (Qsonica, Q800R). To test for DNA-PKcs-S2056p, cells were treated with M3814 or DMSO for 3 hours, treated with 10 Gy IR (MultiRad 160) treatment, and were allowed to recover for 1 hour. Protein was extracted using the ELB solution described above with the addition of PhosSTOP (Roche) and 50 μM sodium fluoride. Blots were probed with antibodies for 53BP1 (Abcam ab36823), RIF1 (Cell Signaling Technologies 95558s), DNA-PKcs (Invitrogen MA5-13238), DNA-PKcs-S2056p (Abcam ab124918), MLH1 (Abcam ab92312), and ACTIN (Sigma A2066). HRP signals were developed using ECL reagent (Amersham Biosciences).

### Clonogenic Survival

To test clonogenic survival, 6-well plates were coated with poly-lysine and cells were plated at various cell densities with 2.5 mL complete media containing M3814 or DMSO. The following day, cells were treated with 1 Gy IR (MultiRad 160) or left untreated. All cells (IR treated and untreated) were treated with 3 mL fresh media containing M3814 or DMSO, and colonies were allowed to form for 7-10 days. Colonies were fixed in cold methanol before staining with 0.5% crystal violet (Sigma) in 25% methanol. Colonies were counted blindly, as in sample identity was hidden from the experimenter conducting the counting, under a 4X objective. Survival was determined for each well relative to the mean value of DMSO/untreated wells for the respective cell line that were seeded in parallel.

## ACKNOWLEDGEMENTS

This study was funded in part by the National Cancer Institute of the National Institutes of Health: R01CA256989 (J.M.S.); P30CA33572 (City of Hope Core Facilities); F99CA284248 (M.C.A.).

## DATA AVAILABILITY STATEMENT

The datasets in the study are included in the study and are available from the corresponding author on reasonable request.

## AUTHOR CONTRIBUTIONS

K.M., M.C.-A., F.W.L, and J.M.S. designed research; K.M., M.C.-A., and F.W.L. performed research; K.M. and J.M.S. analyzed data and wrote the paper with input from all authors.

