## Supplemental Table 1, Supplemental Figure 1-5 for "53BP1/RIF1 and DNA-PKcs show distinct genetic interactions with diverse chromosomal break repair outcomes"

**Supplemental Table S1. Oligonucleotide list.**

| **Name** | **Purpose** | **Sequence (5' → 3', all sgRNA sequences have an initial G nucleotide, regardless of whether it is part of the targeted sequence)** |
|---|---|---|
| 7a (EJ7-GFP-a) | sgRNA | GACCACCCTGACCTACGGCTA |
| 7b (EJ7-GFP-b) | sgRNA | GGCTGAAGCACTGCACGAAT |
| MTAP | sgRNA | GTTACCATGGTCTGTATGGCC |
| CDKN2B-AS1 | sgRNA | GTCGTCACTACGCTGATGGCA |
| LMNA | sgRNA | GCCATGGAGACCCCGTCCCAG |
| 53BP1sg1 | sgRNA | GCATAATTTATCATCCACGTC |
| 53BP1sg2 | sgRNA | GAGAGAATGAGGCTCGAAGTG |
| RIF1sg1 | sgRNA | GAAGTCTCCAACAGCGGCGCG |
| MAfustionHYB | primer | /56-FAM/GAAGGGAGA/ZEN/GAGAGGAGGGA/3IABkFQ/ |
| MTAPctrlDN1 | primer | TCCAGTGTCCTCAAATTCCA |
| MAfusion1UP | primer | AAGTGTGATGGGCAAGAAGG |
| MAfusion1DN | primer | AGGATTCTGCACTTGGATGG |

Supplementary Figure 1

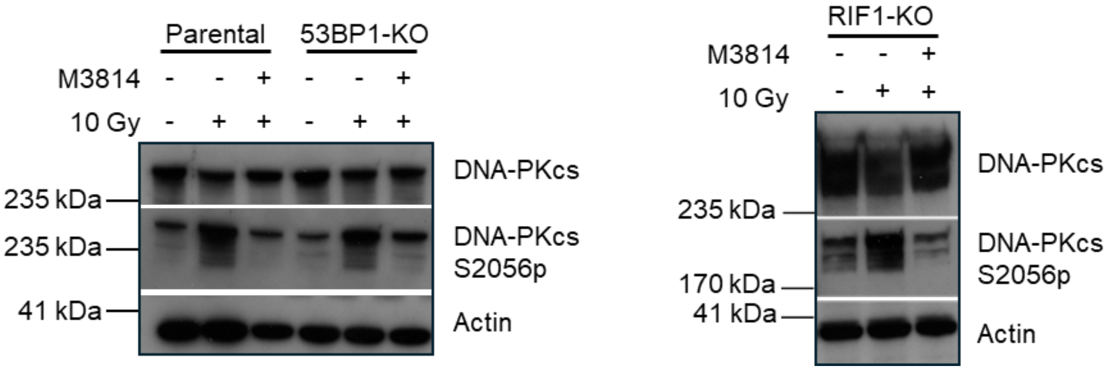


**Supplementary Figure 1. M3814 inhibits DNA-PKcs autophosphorylation independent of 53BP1 and RIF1.** Shown are immunoblot signals for DNA-PKcs, DNA-PKcs S2056p, and Actin from cells (Parental, 53BP1-KO, and RIF1-KO) treated with 500nM M3814 and 10 Gy IR, 10 Gy IR alone, and untreated.

Supplementary Figure 2

A

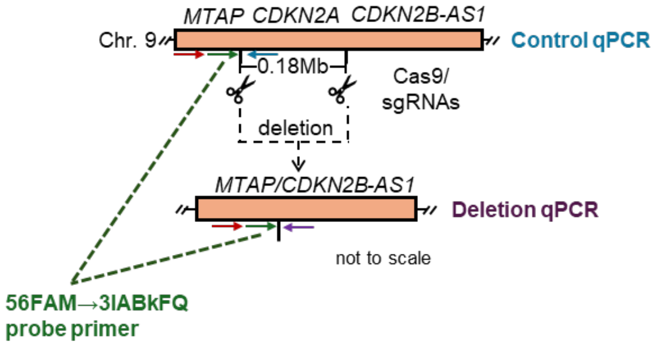


B

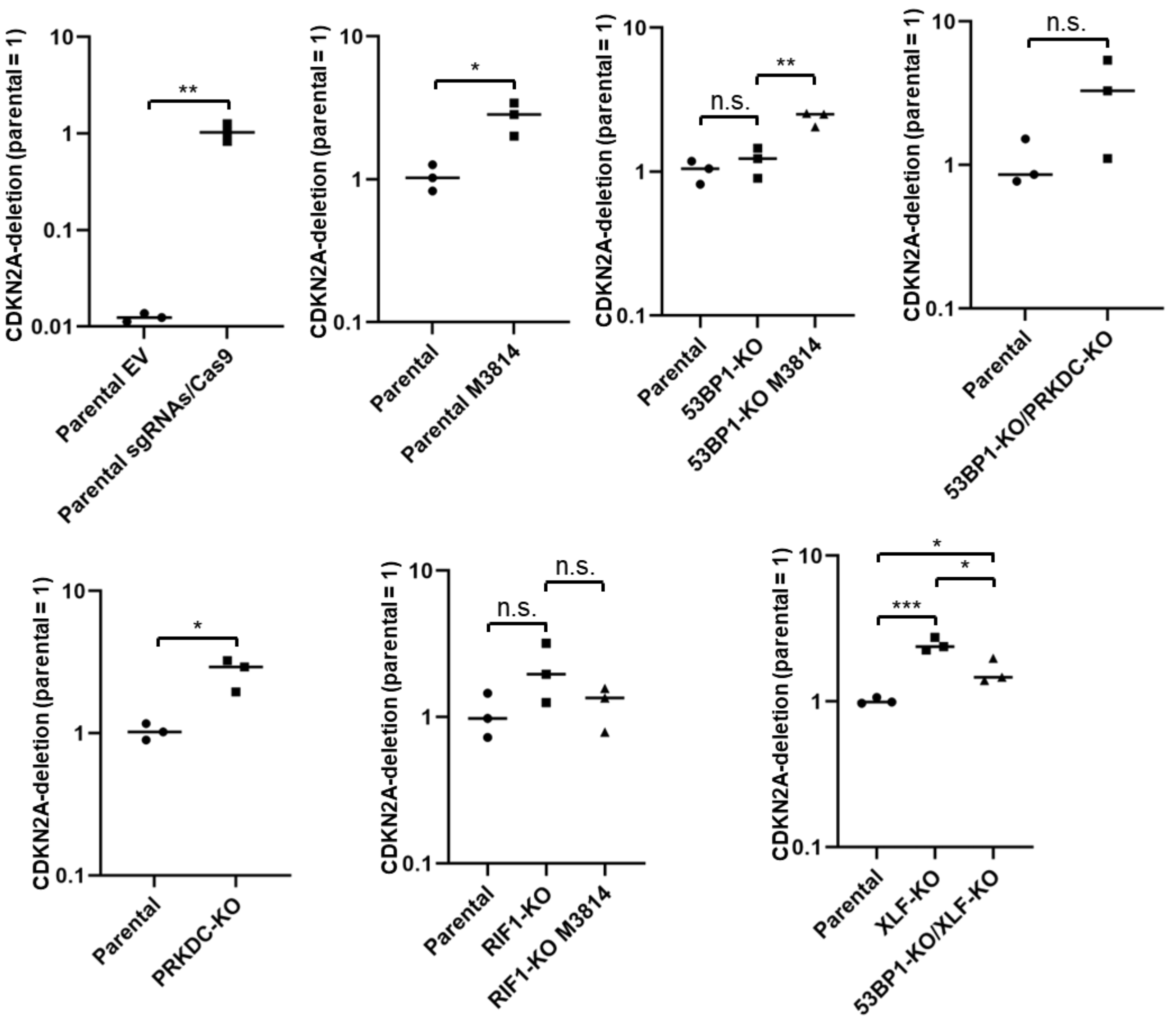

**Supplementary Figure 2. Genetic disruption of 53BP1, PRKDC, RIF1, XLF, and DNA-PKcs kinase inhibition (M3814 treatment) does not cause a decrease in MA-Del deletion frequency.** **(A)** Schematic of the qPCR that involves three primers mapped across the MTAP break locus: the red arrow is the forward primer in *MTAP*, the green arrow denotes the 56FAM→3IABkFQ primer that is the fluorescent probe primer, the blue arrow is the reverse primer for *MTAP* (used for the control reaction), and the purple arrow is the reverse primer in *CDKN2B-AS1* (used to detect the MA-del deletion). **(B)** Deletion frequency is detectable by qPCR with the use of MA-Del Cas9/sgRNAs, and genetic disruption of the factors shown, and M3814 treatment does not obviously cause a decrease in deletion frequency. n=3 biologically independent transfections. Statistics with unpaired t-test using Holm-Sidak correction. *P<0.5, **P<0.01, ***P<0.001, n.s.=not significant.

Supplementary Figure 3

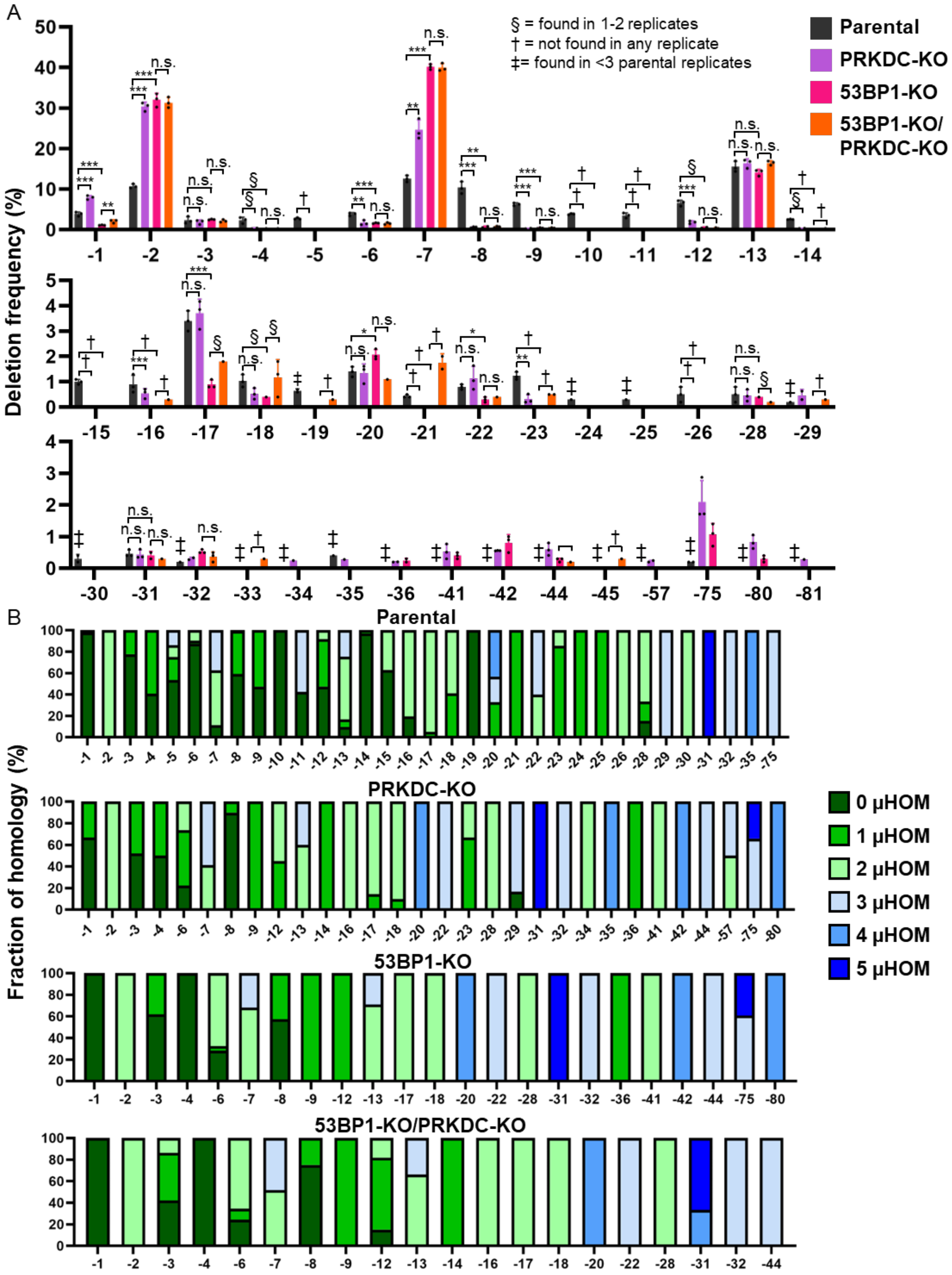

**Supplementary Figure 3. Genetic loss of 53BP1 and DNA-PKcs, alone and together, cause a similar shift in deletion patterns.. (A)** Deletion sizes are similarly effected by genetic loss of 53BP1 and DNA-PKcs (PRKDC), and the combined disruption is similar to the single disruption. n=3 independent transfections. Statistics with unpaired t-test using Holm-Sidak correction. *P<0.05, **P<0.01, ***P<0.001, ****P<0.0001, n.s. = not significant. § = deletion size was only found in 1 or 2 replicates, † = deletion size was not found in any of the replicates, and ‡ = deletion size was found in <2 parental replicates. Parental and 53BP1-KO values are the same as those in Figure 3A. (**B)** Microhomology use varies with distinct deletions. Shown is the fraction of microhomology used for each deletion size for several experimental conditions. n=3 biologically independent transfections. Parental and 53BP1-KO values are the same as those in Figure 3B.

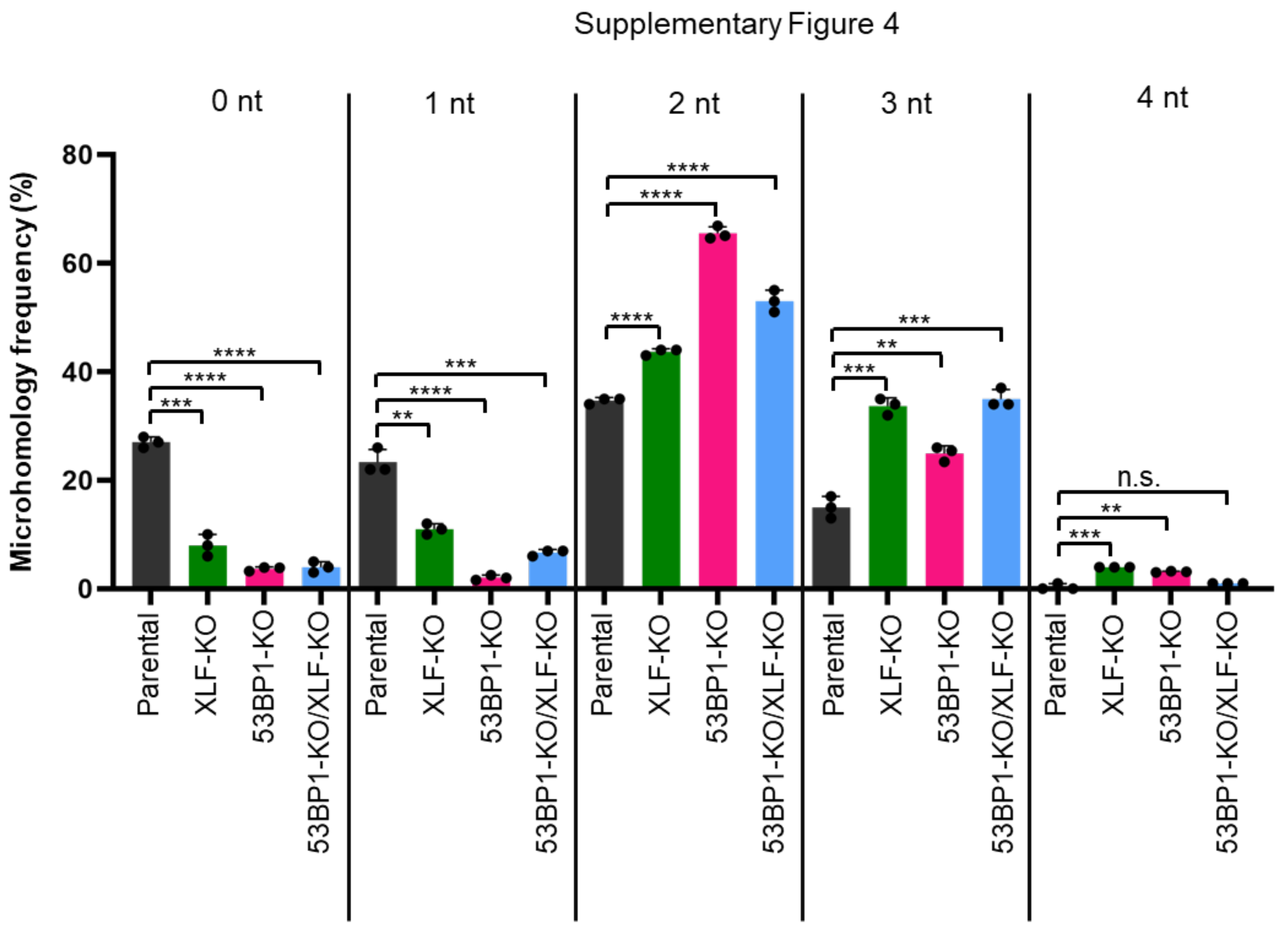


**Supplementary Figure 4. Loss of 53BP1 and XLF, alone and together, cause a similar increase in microhomology usage for deletions.** Shown is the frequency of microhomology used amongst all deletions for each experimental condition. This is microhomology analysis of the data shown in Fig 6. n=3 biologically independent transfections. Statistics with unpaired t-test using Holm-Sidak correction. *P<0.05, **P<0.01, ***P<0.001, ****P<0.0001, n.s. = not significant.

Supplementary Figure 5

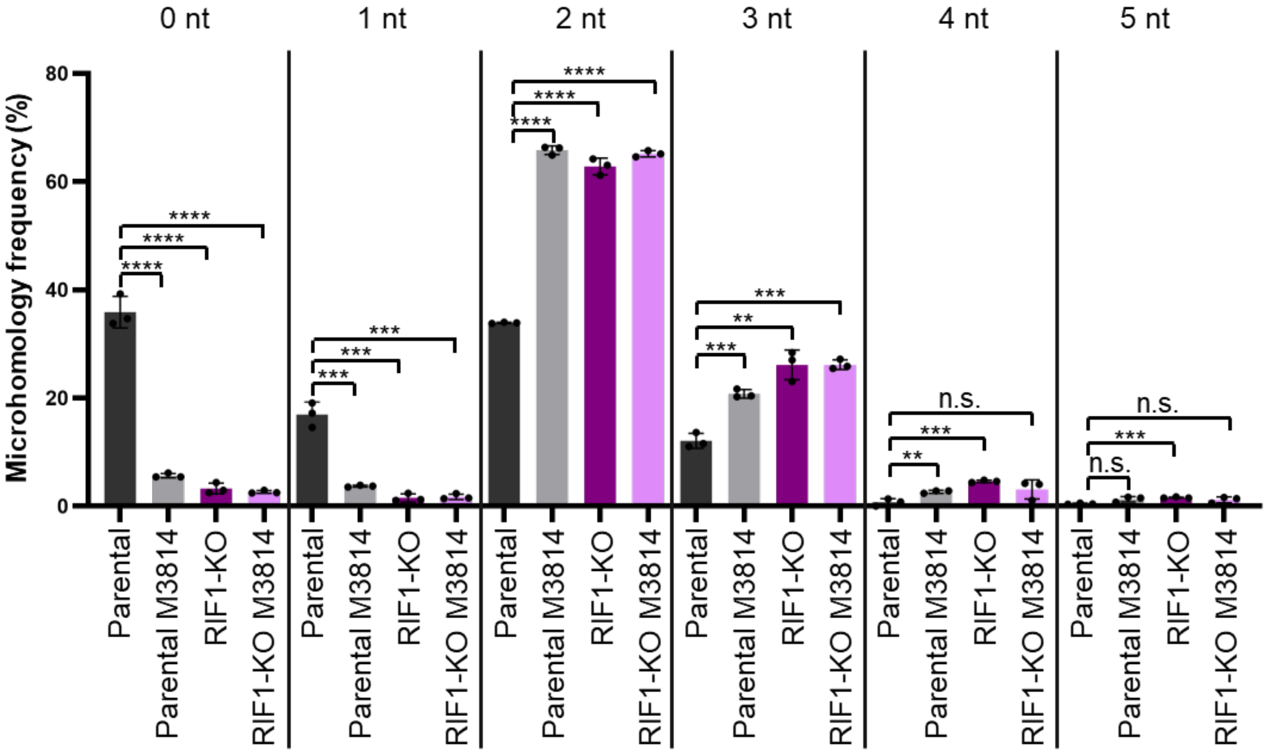


**Supplementary Figure 5. Loss of RIF1 and DNA-PKcs kinase inhibition, alone and together, cause a similar increase in microhomology usage for deletions.** Shown is the frequency of microhomology used amongst all deletions for each experimental condition shown. This is analysis of the data shown in Fig. 8. n=3 biologically independent transfections. Statistics with unpaired t-test using Holm-Sidak correction. *P<0.05, **P<0.01, ***P<0.001, ****P<0.0001, n.s. = not significant.
